# Depletion of endomembrane reservoirs drives phagocytic appetite exhaustion in macrophages

**DOI:** 10.1101/2024.07.31.605905

**Authors:** Aaron Fountain, Mélanie Mansat, Tracy Lackraj, Maria C. Gimenez, Serene Moussaoui, Maya Ezzo, Sierra Soffiaturo, Elijah Urdaneta, Munira B. Verdawala, Karen Fung, Charlene Lancaster, Elliott Somerville, Boris Hinz, Mauricio R. Terebiznik, Roberto J. Botelho

## Abstract

During phagocytosis, a phagocytic cup grows via F-actin remodelling and localized secretion to entrap a particle within a phagosome, which then fuses with endosomes and lysosomes to digest the particle, followed by phagosome resolution. As spatially limited systems, phagocytes have a maximal phagocytic capacity, at which point further uptake must be blunted. However, the processes responsible for phagocytic appetite exhaustion as phagocytes reach their maximal phagocytic capacity are poorly defined. We found that macrophages at their capacity have lower surface levels of Fcγ receptors but overexpression of these receptors did not increase their capacity, suggesting that receptor levels are not limiting. Instead, surface membrane in-folding, membrane tension, and cortical F-actin were all reduced in exhausted macrophages. While this might contribute to appetite suppression, we also found that “free” endosomes and lysosomes were severely depleted in exhausted macrophages. Consequently, focal exocytosis at sites of externally bound particles was blunted. In comparison, macrophages recovered their appetite if phagosome resolution was permitted. We propose that depletion of the endomembrane pools is a major determinant of phagocytic fatigue as macrophages reach their phagocytic capacity.

**Summary statement:** Macrophages that reach their maximal phagocytic capacity lose their appetite for further uptake. This appetite exhaustion is driven partly by depletion of endosomes and lysosomes, preventing growth of additional phagocytic cups.

## Introduction

Phagocytic cells recognize and engulf a variety of particulates that include apoptotic and senescent cells, host-derived cell debris, and potentially pathogenic bacteria, fungi, and protists. These particulates are individually entrapped within a phagosome, an organelle that forms *de novo* using the plasma membrane. Thus, in multicellular organisms, phagocytosis is necessary to clear infections, for antigen processing and presentation, to remodel tissues during development, and to maintain tissue homeostasis and functions (Arandjelovic and Ravichandran, 2015; Flannagan et al., 2012; Fountain et al., 2021; Lancaster et al., 2019; Yin and Heit, 2021). After formation, phagosomes follow a maturation program by fusing with endosomes and lysosomes and thereby converting into highly degradative and acidic phagolysosomes. Phagosome maturation is controlled by various GTPases and lipid signals that coordinate the spatio-temporal sequence of membrane fusion and fission events (Fairn and Grinstein, 2012; Fountain et al., 2021; Hampton and Dickerhof, 2023; Levin et al., 2016; Nguyen and Yates, 2021). The enclosed particulate is thus digested, and its components recycled. After degradation, phagosomes fragment into vesiculo-tubular compartments that recycle membranes and reform lysosomes – this process is referred to as phagosome resolution (Fountain et al., 2021; Lancaster et al., 2021; Levin et al., 2016; Levin-Konigsberg et al., 2019).

At first glance, the primary source of membrane used to form a phagosome is the plasma membrane, which is shaped into a phagocytic cup via F-actin-driven protrusions (Flannagan et al., 2012; Greenberg et al., 1990; Greenberg et al., 1991). Even before binding, plasma membrane extensions probe the cellular environment for and capture potential prey for phagocytosis (Flannagan et al., 2010). However, plasma membrane and actin remodelling do not suffice to form phagocytic pseudopods; exocytosis of endomembranes is required (Masters et al., 2013). For example, recycling endosomes, late endosomes, and Golgi-derived vesicles undergo focal exocytosis at sites of engulfment (Bajno et al., 2000; D’Amico et al., 2021; Hanes et al., 2017; Vashi et al., 2017; Vinet et al., 2008). For larger particles, this also entails secretion of lysosomes (Czibener et al., 2006; Davis et al., 2020; Samie et al., 2013; Sun et al., 2020). Even the endoplasmic reticulum was argued to supply membrane during phagocytosis (Gagnon et al., 2002), though this observation is disputed (Touret et al., 2005). Regardless, the signals that drive localized exocytosis are not well defined but may include increase in membrane tension, actin depolymerization, phosphoinositide signaling, and/or calcium signaling (Bajno et al., 2000; Braun et al., 2004; Czibener et al., 2006; D’Amico et al., 2021; Masters et al., 2013; Samie et al., 2013; Sun et al., 2020). Overall, the incorporation of endomembranes onto phagocytic cups offsets the internalization of plasma membrane during phagocytosis and may even enlarge the phagocyte during phagocytosis (Cannon and Swanson, 1992; Cox et al., 1999; Di et al., 2003; Holevinsky and Nelson, 1998). Taken together, phagocytes seem to consume plasma membrane and/or endomembranes to engulf and internalize particles by phagocytosis.

As spatially limited systems, macrophages must have a maximal phagocytic capacity (heretofore, *phagocytic capacity*), at which point their phagocytic appetite must become exhausted (heretofore, *phagocytic exhaustion or fatigue*). While the mechanisms of phagocytic exhaustion are poorly defined, Zent and Elliott provided an elegant road map to understand possible mechanisms driving phagocytic exhaustion of phagocytes at their phagocytic capacity; this included negative signaling, spatial hindrance, and availability of membrane reservoirs (Zent and Elliott, 2017). However, their relative role in dictating phagocytic exhaustion may depend on phagocyte, particle properties, and phagocytic signaling, and more. For example, membrane availability was proposed to determine the maximum size of a particle that could be engulfed by a macrophage (Cannon and Swanson, 1992) and that a membrane pool participates in phagocytosis (Cannon and Swanson, 1992; Petty et al., 1981). In fact, we previously reported that lysosome number is reduced after phagocytosis of microbeads and filamentous *Legionella*, though we did not explore the fate of endosomes (Lancaster et al., 2021). In comparison, Fcγ receptor (FcγR) levels were reduced in macrophages after extensive phagocytosis of tumour cells opsonized with anti-CD20 antibodies (Pinney et al., 2020), suggesting that decreased receptor levels may drive phagocytic exhaustion. Collectively, multiple mechanisms may regulate phagocytic exhaustion in a manner dependent on phagocytic cell, receptor, and target type.

In this study, we investigated how macrophages exhaust their phagocytic appetite, chiefly using IgG-coated 3 µm beads and RAW macrophages. Under the observed conditions, surface FcγR levels drop in exhausted macrophages, but their overexpression does not increase phagocytic capacity suggesting that receptor levels are not a bottleneck. Moreover, cytoplasmic crowding with phagosomes did not increase mechanical tension in macrophages at their maximal phagocytic capacity; in fact, the plasma membrane appeared less tense and there was a reduction in cortical F-actin. Interestingly, fatigued macrophages were devoid of plasma membrane infolding and depleted of “free” endosomes and lysosomes, suggesting that a loss of membrane reservoirs dictated phagocytic exhaustion in this context. Consistent with this, macrophages recovered their phagocytic appetite upon phagosome resolution to recycle membranes.

## Results

### Macrophages exhaust their phagocytic appetite

Macrophages are spatially limited systems and must have a maximal phagocytic capacity at which point macrophages should stop engulfing additional particles, i.e., their phagocytic appetite is exhausted. Nonetheless, the mechanisms that enable macrophages to sense their phagocytic capacity and accordingly curb their appetite are mostly uncharacterized (Cannon and Swanson, 1992; Zent and Elliott, 2017). It is also not known if and how macrophages recover their appetite after they reach their phagocytic capacity, which likely depends on particle digestion.

To determine their phagocytic limit, we fed RAW264.7 (RAW) macrophages with excess amounts of non-digestible IgG-opsonized 3 µm beads over 4 h. After 120 min of phagocytosis, the phagocytic capacity plateaued indicating that macrophages reached their phagocytic limit at this point and could not engulf additional particles (Fig. 1A, B). We note that the absolute number of phagosomes in RAW macrophages at capacity can vary between experiments and may reflect differences in cell passage, serum lot, or IgG lot. To further ascertain this phagocytic appetite exhaustion, we employed two waves of phagocytosis using different particles. In the first round of phagocytosis, macrophages were kept unfed or presented with 3 µm IgG-coated beads or mRFP1-expressing *E. coli* for 2 h to approach their phagocytic capacity limit. Macrophages were then challenged with a second phagocytic feeding with GFP-expressing *E. coli* for 1 h (Fig. 1C). We observed that naïve macrophages engulfed ∼12 GFP-*E. coli* per macrophage (Fig. 1D, E). By contrast, macrophages that previously internalized IgG-beads or mRFP1-*E. coli* to near capacity (first feeding) consumed significantly fewer GFP-*E. coli* on the second feeding (Fig. 1D, 1E). This also indicates that phagocytic exhaustion occurs within 3 h of uptake whether macrophages engulf digestible or indigestible particles. Thus, based on the current experimental model, macrophages exhaust their phagocytic appetite after 2-3 h of feeding. This phagocytic exhaustion was not due to macrophage damage or metabolic fatigue since control and exhausted macrophages were indistinguishable when stained with Sytox Deep Red, a vital indicator that labels cells if the membrane is damaged, or with Mitotracker red and green, which respectively report on mitochondria membrane potential and mitochondrial number (Sup. Fig. S1).

**Figure 1:**
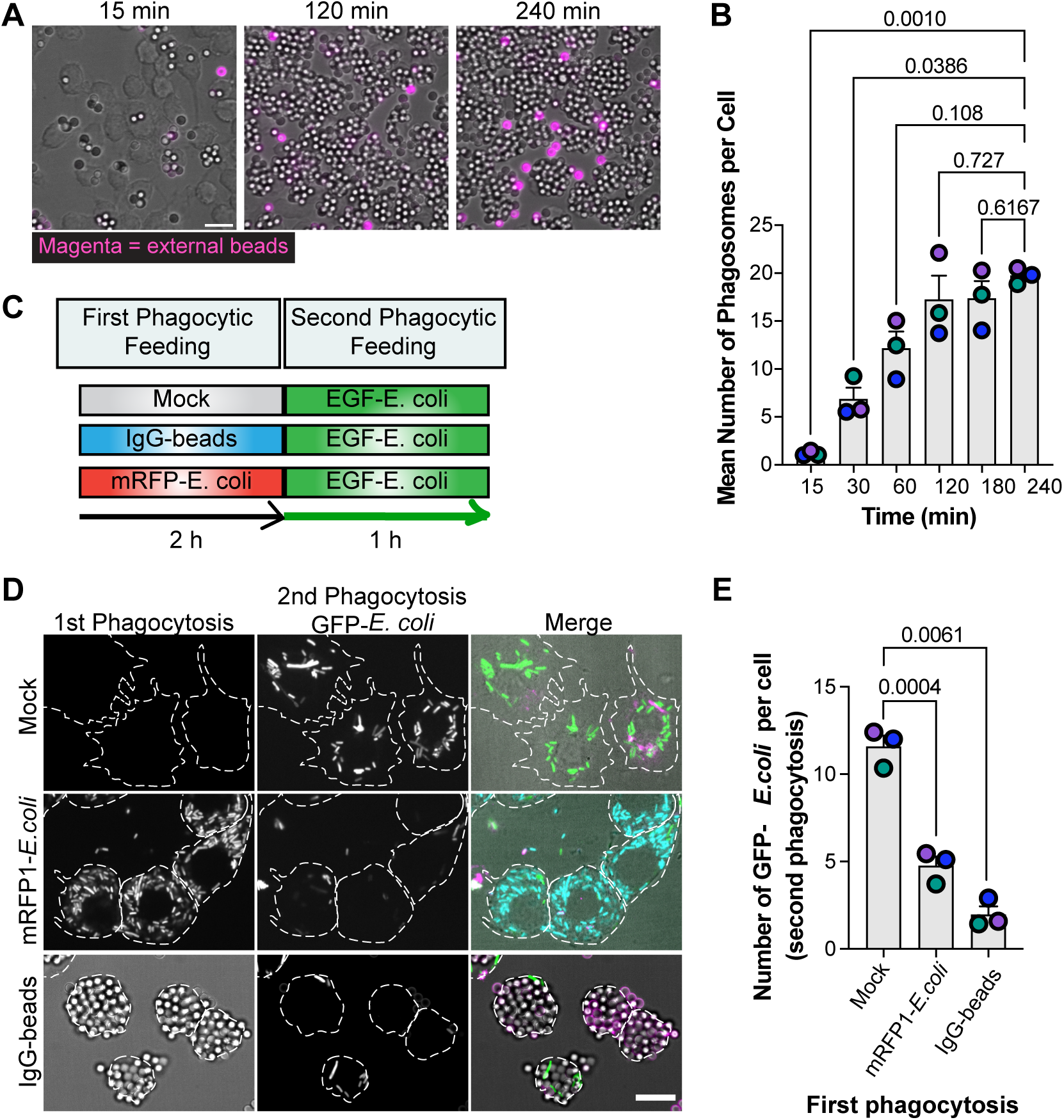
Macrophages curb their phagocytic appetite after two hours of engulfment. **A.** RAW macrophages were allowed to engulf IgG-opsonized 3 µm beads for 0 to 240 minutes (15, 120, and 240 min shown). External particles were immunolabeled after fixation with anti-human IgG secondary antibodies (magenta). **B.** Number of internalized beads per cell as described in A. Data is shown as mean ± SEM from 3 independent replicates, where 200-300 cells per condition per experiment were quantified. Sample means were compared against the 240 min timepoint. **C.** Schematic of sequential phagocytic feeding assays to test loss of phagocytic appetite in macrophages. Macrophages were subjected to a first phagocytic feeding for 2 h (mock, IgG-beads, mRFP1-*E. coli*), followed by an immediate second feeding with GFP-*E. coli* for 1 h. **D**. RAW cells were pre-fed mRFP1-*E. coli* (shown as cyan in Merge channel), or IgG-beads, or were mock fed, followed by a second round of phagocytosis using GFP-*E. coli* as described in C. Magenta indicates external particles. For A and D, dashed lines delineate cell boundaries. Scale bar: 10 µm **E.** Number of GFP-*E. coli* engulfed by RAW cells as described in D. Data are shown as mean ± SEM from 3 independent replicates, where 100-200 cells per condition per experiment were quantified. For B and E, data was tested using a repeated-measures one-way ANOVA and Dunnett’s post-hoc test, where p-values are shown.

### Total and surface levels of Fcγ receptors remain unaltered in exhausted macrophages

Various processes may drive macrophage phagocytic exhaustion, including depletion of phagocytic receptors, which was observed during antibody-dependent phagocytosis of tumour cells (Pinney et al., 2020). Since FcγRIIIA (CD16A) and FcγRI (CD64) are thought to be the main phagocytic receptors for IgG-coated particles in RAW macrophages (Hayes et al., 2016; Willcocks et al., 2009), we measured their surface levels after phagocytosis of IgG-beads over 2 h. First, we observed a gradual decrease in anti-CD16A antibody fluorescence by microscopy (Fig. 2A-B) and by flow cytometry (Fig. 2C, Sup. Fig. S2A), despite the total levels remaining the same (Fig. 2E-G). We also observed a decrease in anti-CD64 fluorescence by flow cytometry (Fig. 2C, Sup. Fig. S2B; we avoided microscopy of CD64 because indirect immunofluorescence caused CD64 clustering). Yet, macrophages at capacity were often decorated by externally bound beads (Fig. 2A, insets; Fig. 2H), which might sterically block anti-CD16A and anti-CD64 antibodies from accessing respective surface receptors.

**Figure 2:**
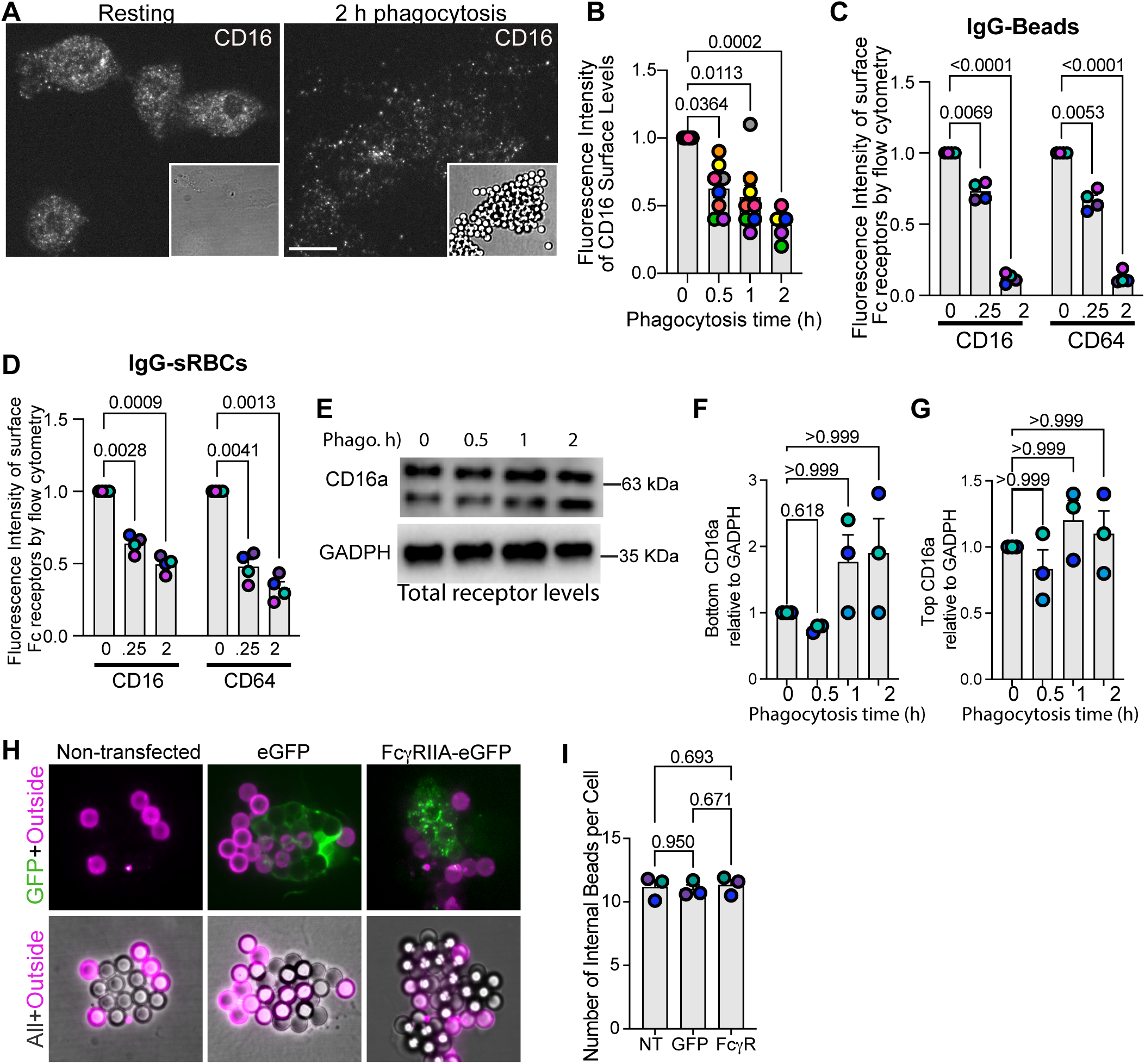
Surface Fcγ receptor levels are reduced in exhausted macrophages, but this does not drive appetite exhaustion. **A.** RAW cells were fed IgG-coated 3 µm beads for the indicated times, followed by fixation, and staining with anti-CD16A. Cells were imaged along the z-axis and reconstructed by summing the fluorescence intensity. Scale bar = 10 µm. Inset shows corresponding bright field showing beads. **B:** Normalized CD16A mean fluorescence intensity of reconstructed z-stacks per cell from n=5-7 independent experiments from 50-100 cells per experiment per condition. **C, D:** RAW cells were fed IgG-coated beads (C) or IgG-coated sRBCs (D) for indicated times and then processed for flow cytometry to stain against CD16A or CD64. Shown is the mean fluorescence for each receptor normalized to resting cells (0 min; Supplemental Figure S2 shows representative flow cytometry distribution). **E-G.** Western blotting of whole cell lysates of macrophages fed IgG-beads for indicated times (E) and the quantification of bottom bottom (F) and top (G) CD16A bands relative to GADPH. **H.** RAW cells were mock transfected or transfected with plasmids encoding eGFP or FcγRIIA-GFP (green) and then allowed to phagocytose 3 µm IgG-beads for 2 h, followed by fixation and staining of external beads (magenta). Scale bar = 10 µm. **I.** Number of internalized beads per cell from n=3 independent experiments, counting 50-100 cells per condition per experiment. B: Data was quantified by Kruskal-Wallis and Dunn’s post-hoc test. C, D: Repeated-measures two-way ANOVA followed by Tukey’s post-hoc test. F, G: Data were analysed by Friedmann’s test, followed by Dunn’s post-hoc test. I. Data was analysed by One-way ANOVA and Tukey’s post-hoc test. All p values shown.

To allay this concern, we fed RAW cells IgG-coated sheep red blood cells (sRBCs); externally bound sRBCs can be lysed by hypoosmotic shock to form membrane ghosts that we reasoned would permit antibody-receptor binding. Using flow cytometry, we still observed a reduction in surface levels of CD16A and CD64 in macrophages that engulfed IgG-sRBCs to capacity (Fig. 2D, Sup. Fig. S2C, S2D).

These observations imply that surface FcγR levels may be limiting and drive phagocytic exhaustion as macrophages approach their phagocytic capacity. If so, over-expression Fcγ receptors should increase the phagocytic capacity of macrophages. Nonetheless, over-expression of FcγRIIA-GFP in RAW cells did not enhance their phagocytic load after 2 h of particle uptake (Fig. 2H, I). Collectively, while surface FcγR levels may be lower in exhausted macrophages, this does not seem to be a limiting factor that authorises phagocytic exhaustion as macrophages approach their capacity, suggesting that other processes are at play.

### Cell volume and surface area of exhausted macrophages are not reduced

Prior reports indicate that post-phagocytosis, macrophages lose their plasma membrane infolding, and either retain or enlarge their cellular volume and surface area (Cannon and Swanson, 1992; Petty et al., 1981). However, these studies may not have considered macrophages at their maximal phagocytic load. Conceivably, a macrophage that embraces as many particles as possible may suffer a reduction in surface area and/or volume as they consume membrane to form phagosomes. In turn, such a reduction in cell size could then curb phagocytic appetite due to physical constraints. We thus assessed membrane infolding and cell surface area and volume in exhausted macrophages.

The macrophage membrane surface is enriched in membrane infolds, filopodia, lamellipodia, and other extensions (Araki et al., 1996; Condon et al., 2018; Petty et al., 1981). To resolve the extent of membrane infolding of the plasma membrane in resting and exhausted macrophages, we turned to scanning electron microscopy. Indeed, resting macrophages were remarkably enriched in membrane folds, sheets, and various forms of extensions (Fig. 3A, B). In comparison, macrophages that engulfed 3-µm beads for 15 min retained a complex, convoluted membrane morphology with some beads appearing to “bulge out” from within (Fig. 3A). However, macrophages at capacity displayed a striking “raspberry” morphology, with internalized beads appearing to distend the macrophage surface, while the surface was divested of membrane folds, being mostly smooth or “bald” (Fig. 3A, B, and D). This implies that exhausted macrophages have depleted all their surface “give” to accommodate as many particles as possible, an observation congruent with (Petty et al., 1981).

**Figure 3:**
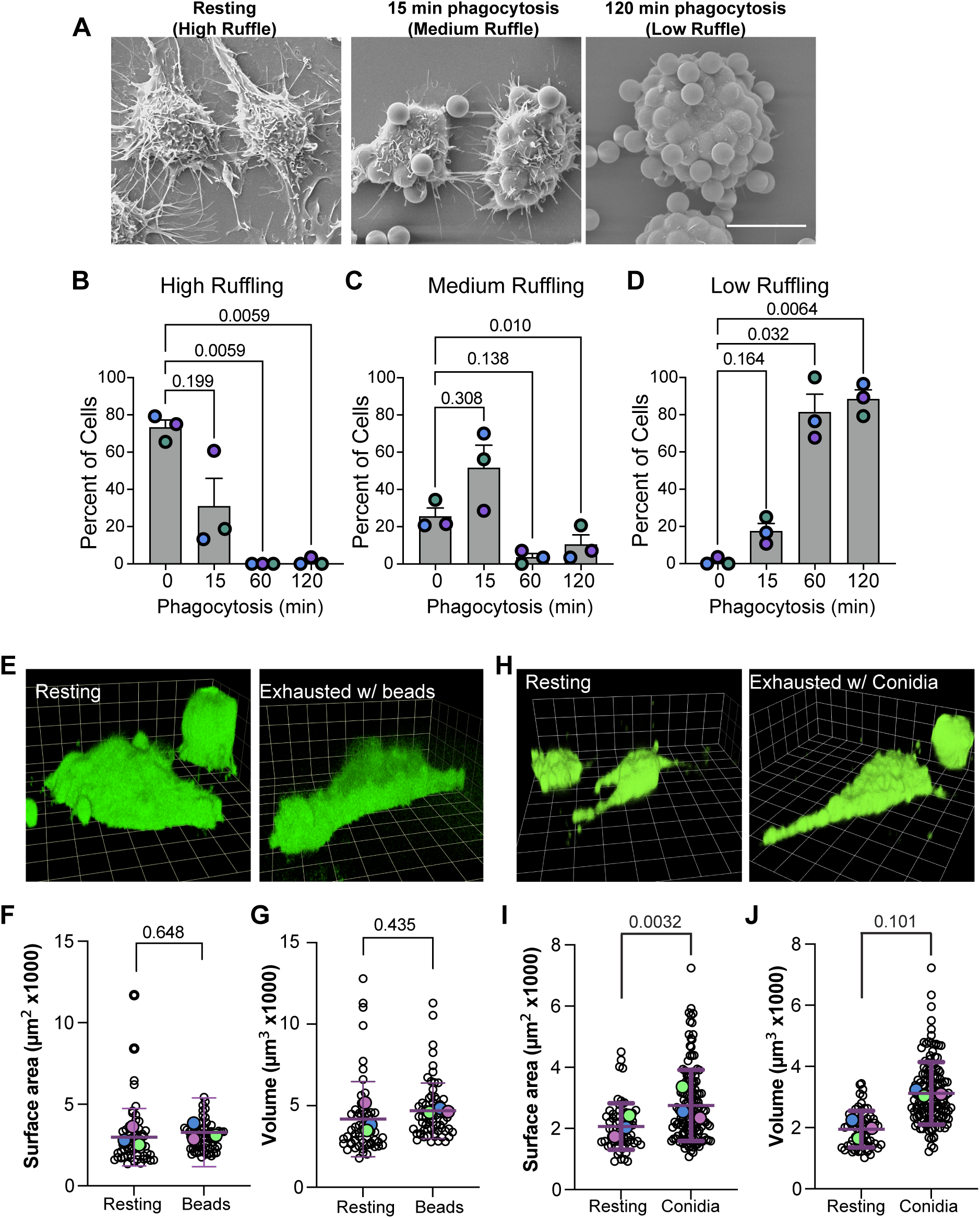
Macrophages become devoid of plasma membrane infolding but do not shrink after phagocytic exhaustion. **A.** Representative images acquired by SEM of the plasma membrane ultrastructure of macrophages after 0, 15, or 120 min of phagocytosis of IgG-opsonized beads; left panel: resting macrophages have a high degree of membrane ruffling (High Ruffle) with many large membrane sheets and protrusions emanating from the cell surface, creating valleys and pits on the cell surface; middle panel: macrophages that engulfed a few particles have reduced, but visible membrane ruffling (Medium Ruffle) across at least 50% of the apical surface of the cell; right panel: macrophages at phagocytic capacity are mostly devoid of membrane ruffles (Low Ruffle) with most being extremely shallow. Instead, cells appear as a “raspberry” due to engulfed beads protruding from the cytosol-out. Scale bar: 10 µm. **B, C**, and **D**. Quantification of percent cell population exhibiting high (B), medium (C), or low (D) degree of ruffling as described in A. Data represented as mean ± SEM of 3 independent experiments and from 25-35 cells per replicate per condition. Sample means were analyzed using repeated measures One-way ANOVA and Dunnett’s post-hoc test by comparing to resting cells. **E, H.** Macrophages were transfected to express eGFP-CAAX, a plasma membrane maker prior to phagocytosis. Macrophages were then given no particles or allowed to engulf IgG-opsonized beads (E) or yeast conidia (H) for 2 hours. Images of resting cells (left) and cells containing at least 5 internalized beads or conidia (right) were volumetrically reconstructed from image stacks of EGFP-CAAX fluorescence. Grid lines represent 4.7 µm intervals. **F, G, I,** and **J**. Quantification of surface area (F, G), and volume (I, J) of 15-25 cells per sample as described in *E* and *H*. Data represented as mean ± SEM of 3 independent replicates. White circles represent individual cells, while coloured circles are sample means, which were compared using a paired, two-tailed Student’s t-test.

We then tested whether this corresponded to a decrease in cell volume and/or surface area by decorating the plasma membrane with GFP-CAAX (Madugula and Lu, 2016). Using imaging volumetrics of GFP-CAAX-expressing RAW macrophages, we observed that macrophages that engulfed IgG-beads to capacity did not change their cell volume and surface area relative to resting macrophages (Fig. 3E-G), but those that entrapped conidia increased their surface area (Fig. 3H, I) and tended to become larger (Fig. 3J). This disparity between IgG-beads and yeast conidia uptake may reflect biological differences or may be due to bead-induced light refraction. Regardless, both models indicate that macrophages reach phagocytic capacity without sacrificing their cellular volume and gross surface area.

### The plasma membrane of exhausted macrophages is not under increased tension and is divested of cortical F-actin

The loss of plasma membrane folds and “raspberry” morphology of exhausted macrophages could indicate that the plasma membrane is under higher turgor pressure due to phagosome crowding of the cytoplasm, which in turn, could thwart remodelling of the plasma membrane to engulf additional particles. To appraise the plasma membrane tension, we used fluorescence lifetime imaging microscopy (FLIM) of the Flipper^®^ membrane probe which is sensitive to lipid crowding, an indicator of tension and/or lipid composition (Chen et al., 2023). After the respective treatment, macrophages were labelled with Flipper^®^ and imaged live. Interestingly, macrophages at capacity exhibited reduced lifetime of FLIPPER^®^, which indicates lower lipid crowding and potentially lower tension (Fig. 4A, B). By comparison, the lifetime of FLIPPER^®^ was higher on macrophages exposed to a hypotonic shock, indicating higher membrane tension (Fig. 4A, B).

**Figure 4:**
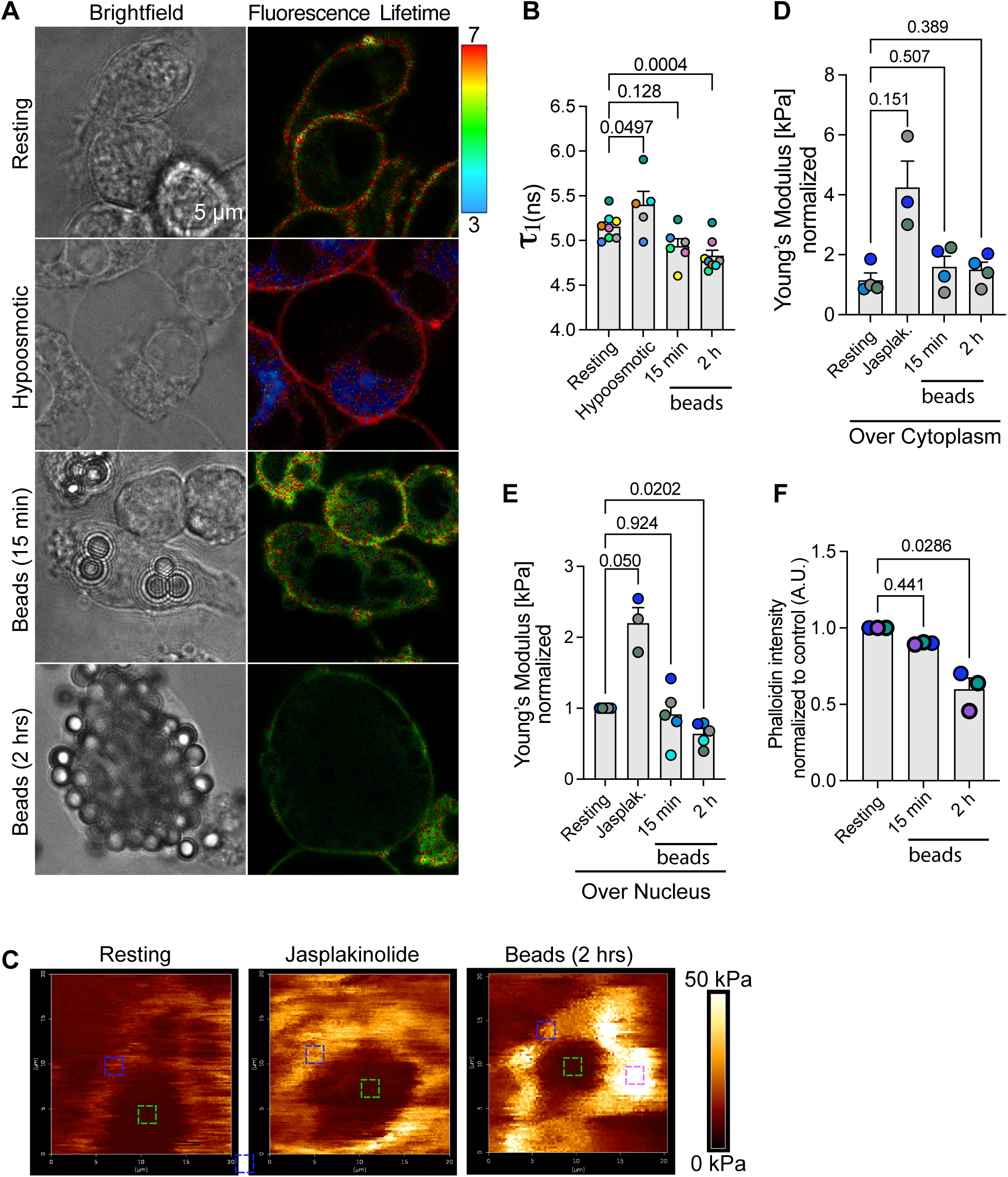
The plasma membrane of exhausted macrophages is not under increased tension. **A.** RAW macrophages were allowed to engulf IgG-opsonized 3 µm beads for 0, 15 min or 2 h, or treated with a hypo-osmotic medium. The cells were then labeled with the Flipper^®^ membrane tension probe and the fluorescence lifetime τ1 was assessed. Brightfield and fluorescence time are shown, where lifetime is pseudo-coloured according to the colour palette scale. **B.** Quantification of the fluorescence lifetime τ1 described in A. Data are the mean ± SEM from at least 5 independent replicates and based on 10-30 cells per condition per experiment. Sample means were compared using mixed-effect one-way ANOVA and Dunnett’s post-hoc test. p-values are shown. **C.** RAW macrophages engulfed IgG-opsonized 3 µm beads for 0, 15 min or 2 hours, or were treated with 1 µM jasplakinolide for 1 h. Illustrative images of cells generated by QI mode shows spatial tension in kPa as indicated by the pseudo-colour palette scale. Intense white signal are beads on the cell. Coloured dashed squares represent areas of measurements, excluding beads. **D, E.** Quantification of the cellular Young’s elastic modulus in kilopascals (kPa) over the cytoplasmic area (D) and over nuclear region (E). Data are shown as mean ± SEM from at least 3-5 independent replicates with the exact repeats represented as individual data points. Per experiment and per condition, 4-6 cells were evaluated at two nuclear positions and four cytosolic positions, each generating 20 curves per position. Data were compared by mixed-effect one-way ANOVA and Dunnett’s post-hoc test with p-values disclosed. **F.** Quantification of whole phalloidin fluorescence intensity normalized to resting macrophages. Data are shown as mean ± SEM from N=3 independent experiments using >40 cells per experiment per treatment. Data was analysed by Friedman test and Dunn’s post-hoc test. p values are disclosed.

Given that lipid crowding is impacted by both membrane tension and lipid composition, we next employed Atomic Force Microscopy (AFM) to better understand these observations. Using Quantitative Imaging (QI), we could generate force-based images as shown in Fig. 4C, which displayed areas of low tension (typically, plasma membrane over the nucleus) and regions that were over 50 kPa (mapped onto beads and phagosomes protruding onto the plasma membrane). While the AFM probe is fine enough to evade areas of the plasma membrane with beads and protruding phagosomes, we opted to separately measure the tension of the plasma membrane atop the nucleus and of the plasma membrane over other regions since the former was mostly devoid of beads, while the latter may still be affected by proximal phagosomes and beads (*i.e.,* the raspberry morphology in Fig. 3).

Interestingly, while we did not observe a significant change in the average Young’s Elastic Modulus over cytosolic regions relative to resting cells (Fig. 4D), there was a significant drop in membrane tension over the nucleus in macrophages at phagocytic capacity relative to resting cells (Fig. 4E). Together with the Flipper® data, we interpret these observations to mean that the surface of exhausted macrophages is not under increased tension; if anything, tension is reduced. Consistent with this notion, phalloidin fluorescence was abated in exhausted macrophages relative to resting macrophages or those with only a few phagosomes (Fig. 4F; also see Fig. 6). In comparison, macrophages treated with jasplakinolide to stabilize the actin cortex displayed higher surface tension over the cytosolic and nuclear regions relative to resting cells (Fig. 4D, E). Overall, we suggest that cytoplasmic crowding does not cause tension to prevent further phagocytosis in macrophages at capacity.

### Free endosome and lysosome populations are depleted in exhausted macrophages

Our evidence so far suggests that phagocytic appetite exhaustion in macrophages at capacity is not likely due to cell shrinkage or cytoplasmic crowding leading to membrane distention. Given that nascent phagosomes are born through local secretion of endosomes and lysosomes at the nascent phagocytic cups (Bajno et al., 2000; Czibener et al., 2006; Lee et al., 2007; Samie et al., 2013; Sun et al., 2020), we next postulated if depletion of membrane reservoirs could curb appetite in macrophages at their phagocytic capacity. To test this hypothesis, we first quantified free lysosomes and endosomes in naïve and exhausted macrophages by fluorescence microscopy, electron microscopy, and membrane fractionation.

We previously showed by fluorescence imaging that “free” lysosomes were consumed during phagosome maturation (Lancaster et al., 2021). We corroborate these observations here via two approaches. First, we again estimated and observed that the number of LAMP1-labelled puncta, which we define here as “free lysosomes”, declined in macrophages that engulfed long filamentous *Legionella* (PFA-fixed) bacteria relative to resting cells (Fig. 5A, B). Secondly, to distinctly assess the loss of “free” lysosomes in exhausted macrophages, we separated phagosomes containing magnetic beads from the residual membrane pool after cell lysis and quantified the relative LAMP1 signal to GADPH signal. As expected, LAMP1 was enriched in the phagosomal fraction, while the mock phagocytosis sample had no LAMP1 (Fig. 5C-E). Notably, we observed a significant loss of LAMP1 to GADPH in macrophages exhausted for phagocytosis versus naïve macrophages (Fig. 5C-E), demonstrating that macrophages at maximal phagocytic capacity are drained in their complement of “free” lysosomes.

**Figure 5:**
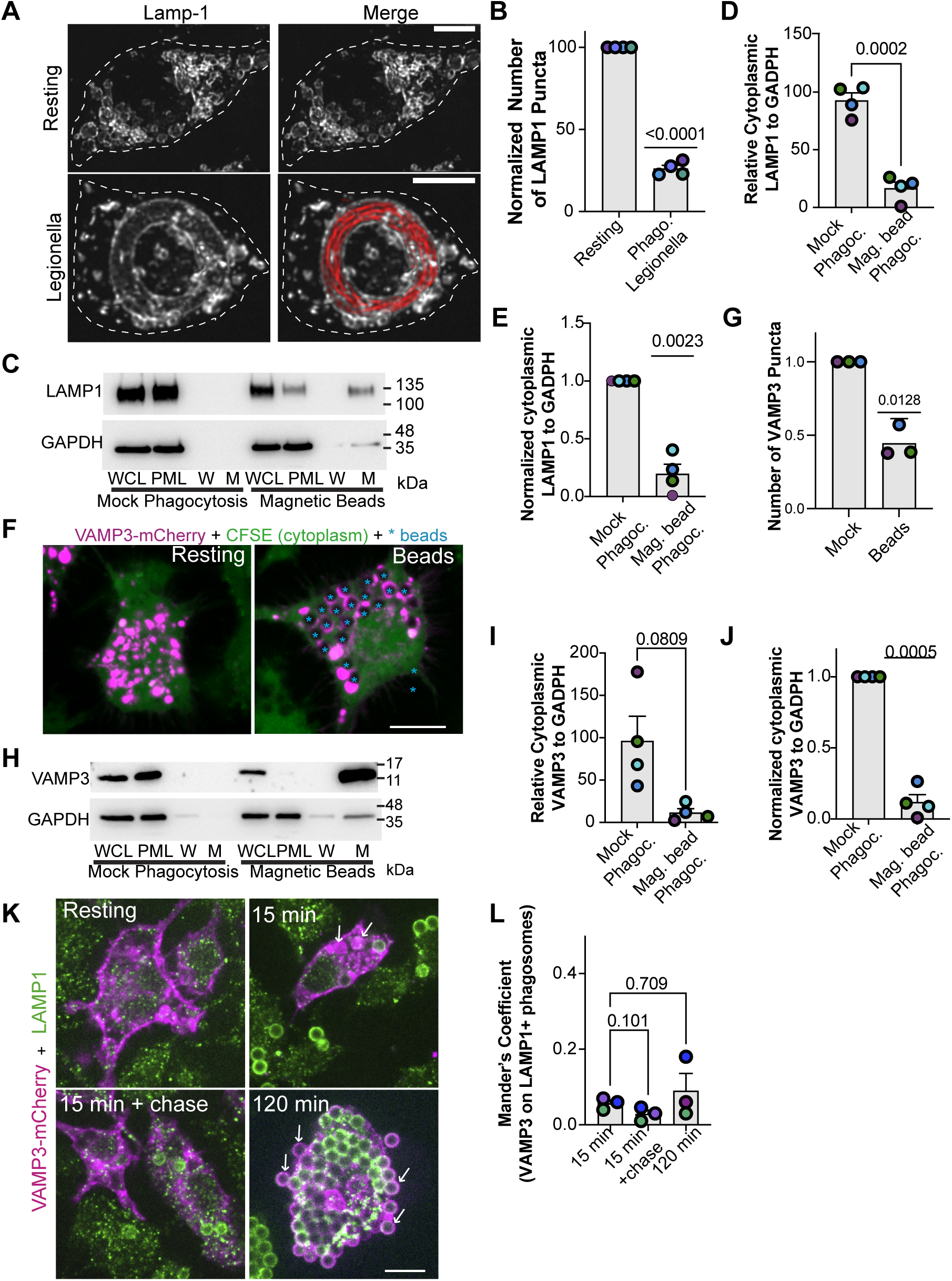
“Free” lysosomes and endosomes are depleted in macrophages at phagocytic capacity. **A.** Reconstructed micrographs of LAMP1 staining (grayscale) in resting and macrophages that engulfed filamentous *Legionella* (red). Phagosome-associated LAMP1 was defined as LAMP1 that colocalized onto and to the periphery of *Legionella*. Dashed lines contours the cell. Free lysosomes were interpreted as LAMP1 puncta not proximal to phagosomes. Scale bar = 10 µm. **B.** Quantification of LAMP1-puncta per cell as described in A. **C, H.** Macrophages were given no particles (Mock phagocytosis, lanes 1-4) or allowed to engulf BSA-anti-BSA-opsonized magnetic beads (Magnetic beads, lanes 5-8) for 2 h, followed by 1 h incubation to elicit maturation. Phagosomes were then magnetically isolated and analyzed with Western Blot. LAMP1 (C) and VAMP3 (H) abundance were probed as a proxy for lysosome and endosome content, respectively. GAPDH was used as a loading control in whole cell lysates (WCL). Post-magnetic lysate (PML) is content remaining in lysate after magnetic separation; Wash fractions (W); Magnetic phagosome fraction is represented by M. **D, I**: Quantification of percent LAMP1 (D) and VAMP3 (I) remaining in lysate after magnetic separation. Percent remaining protein is defined as the percent ratio of GAPDH-normalized protein content in the PML fraction (lanes 2 and 6 for mock phagocytosis and magnetic beads, respectively) to GAPDH-normalized protein content in the WCL fraction (lanes 1 and 5 for mock phagocytosis and magnetic beads, respectively). **E, J**: Analysis in C and H, respectively, but normalized to mock phagocytosis to account for experimental variation in absolute values. Data represented as mean ± STD of N=4 independent experiments. Sample means were compared using two-tailed, paired Student’s t-test. *: p<0.05; **: p<0.01 (D, I) or one sample t and Wilcoxon test (E, J). **F.** Macrophages were transfected to express VAMP3-GFP (magenta) and then were given no particles (resting) or allowed to engulf IgG-opsonized beads for 2 h. Cytosol was demarcated with CFSE staining (green). Phagosome-associated VAMP3 was defined as VAMP3 that colocalized to the phagosome periphery as defined by black void in CFSE resulting from the bead contour. Free endosomes were interpreted as small, VAMP3 puncta not proximal to phagosomes. Scale bar = 10 µm. **G.** Quantification of VAMP3-puncta per cell as described in E. Data represented as mean ± SEM of N=3 independent experiments using 24-40 individual cells per condition per replicate. Data was analysed two-tailed, paired Student’s t-test. **K.** RAW macrophages expressing VAMP3-mCherry (magenta) engulfed IgG-opsonized 3 µm beads for 15 min, with or without 60 min chase, or for 120 min. LAMP1 was immunolabeled after fixation (green). Scale bar = 10 µm. **L.** Manders’ Coefficient of VAMP3-mCherry on LAMP1 phagosomes. Data represented as mean ± SEM of N=3 independent experiments using 30-50 individual cells per condition per replicate. Data was analyzed using repeated-measures one-way ANOVA and Dunnett’s post-hoc test. p-values are shown.

We then evaluated whether exhausted macrophages were also depleted for early/recycling endosomes by probing for VAMP3, a membrane intrinsic SNARE protein (Bajno et al., 2000; Vinet et al., 2008). While phagosomes are expected to become lysosome-like during maturation, and hence use up lysosomes, the early endosome-stage of phagosome maturation is transient (Fountain et al., 2021; Vieira et al., 2003). Thus, *a priori*, one may anticipate that macrophages reform and maintain a free endosome population even after phagocytic exhaustion. Nevertheless, our data suggest that this is not the case. First, we observed a loss of free VAMP3-labelled puncta by fluorescence microscopy in macrophages that engulfed IgG-coated beads to capacity (Fig. 5F, G) or after engulfing long filamentous *Legionella* (Supplemental Fig. S3A, qualitative analysis). Second, we revealed a large depletion of VAMP3 from the residual membrane fraction relative to GADPH in exhausted macrophages compared to naïve macrophages. Instead, VAMP3 was mostly associated with the phagosomal fraction (Fig. 5H-J). Consistent with depletion of endosomes and lysosomes in exhausted macrophages, we observed a decline in endosome/lysosome-like organelles in macrophages that extensively engulfed yeast conidia by transmission electron microscopy (Supplemental Fig. S3B, C). We also observed depletion of “free” LAMP1 and VAMP3 in bone-marrow derived macrophages that engulfed IgG-coated magnetic beads, showing that this is not specific to RAW macrophages (Supplemental Fig. S3D-H). Since both lysosomes (LAMP1) and endosomes (VAMP3) were depleted in exhausted macrophages, we tested if some phagosomes displayed a hybrid endosome-lysosome maturation state by microscopy. However, as illustrated in Fig. 5K-L, phagosomes did not exhibit increased co-localization of VAMP3 and LAMP1 in exhausted macrophages vs. macrophages with a few phagosomes. This implies that phagosomes face a maturation “traffic jam” in exhausted macrophages and that phagolysosomes must be resolved for younger phagosomes to continue maturing.

### Exhausted macrophages cannot secrete endosomes to grow phagocytic cups

We next postulated that exhausted macrophages would not secrete endosomes onto forming cups due to endosome depletion, resulting in shallow phagocytic cups. Indeed, while naïve macrophages readily accumulated VAMP3 on nascent phagocytic cups, exhausted macrophages with externally bound particles were devoid of VAMP3 despite similar levels of total VAMP3-mCherry between resting and exhausted macrophages (Fig. 6A-C). F-actin staining under these externally bound particles was also feeble (Fig. 6A), and in fact, we observed reduced total F-actin staining in exhausted macrophages (Fig. 6D), consistent with observations depicted in Fig. 4F. Moreover, relative to the resting counterparts, we also observed a dampening of total F-actin staining in bone-marrow derived macrophages after engulfing IgG-coated beads or IgG-sRBCs to capacity irrespective of biological sex of the mice (Sup. Fig. S4). Overall, our data indicate that macrophages at their phagocytic limit are depleted of free endosomes and lysosomes, entrapping these membranes as phagosomes. Without this membrane reservoir, we propose that macrophages are unable to grow phagocytic cups and engulf additional particles, triggering phagocytic exhaustion.

### Phagosome resolution recovers the macrophage phagocytic appetite

Our observations suggests that deprivation of membrane reservoirs like endosomes and lysosomes, which are needed for complete nascent phagosomes, drives phagocytic appetite exhaustion of macrophages at capacity. We thus anticipated that phagosome resolution would recover phagocytic appetite by reforming membrane reservoirs. To evaluate this, we again employed two rounds of phagocytosis whereby macrophages either remained naïve, or were fed digestible mRFP1-*E. coli*, or consumed indigestible IgG-coated beads that are incapable of phagosome resolution. These macrophage populations were then chased for 0 h (no resolution) or for 6 h to attempt phagosome resolution (Fig. 7A shows schematic). Macrophages engulfed a high number of mRFP1-*E. coli* (Fig. 7B, C) and beads (Fig. 7B, D) during the first feeding. After 6 h chase, most mRPF1*-E. coli* phagosomes were digested (Fig. 7B, C), while macrophages retained a similar number of beads, being non-digestible, after the first hour of engulfment or after 6 h chase (Fig. 7B, D). After the specific chase time, macrophages were then served GFP-*E. coli* for 1 h as a second feeding to measure their remaining appetite. We disclose the absolute number (Fig. 7E) and normalized number of engulfed GFP-*E. coli* relative to mock phagocytosis (Fig. 7F) to account for experimental variability and because the absolute number of phagosomes in a macrophage at capacity can vary. As illustrated before, macrophages that previously phagocytosed mRFP1-*E. coli* or beads and were allotted no chase period before the second feeding (0 h), had diminished phagocytosis of GFP-*E. coli* relative to macrophages that were kept naïve (Fig. 7B, E, F). In comparison, cells that were allocated 6 h to resolve mRFP1-*E. coli* phagosomes recovered their appetite for GFP-*E. coli* and may even engulf more than naïve macrophages (Fig. 7B, E, F). By contrast, macrophages that first engulfed beads displayed similarly low levels of GFP*-E. coli* phagosomes whether they were given 0 h or 6 h to recover, denoting that phagosome resolution is necessary for macrophages to recuperate their phagocytic appetite (Fig. 7B, E, F). Moreover, the digestibility of and the rate of particle digestion may affect the overall kinetics and dynamics of phagocytic exhaustion and recovery once phagosome resolution ensues.

**Figure 6:**
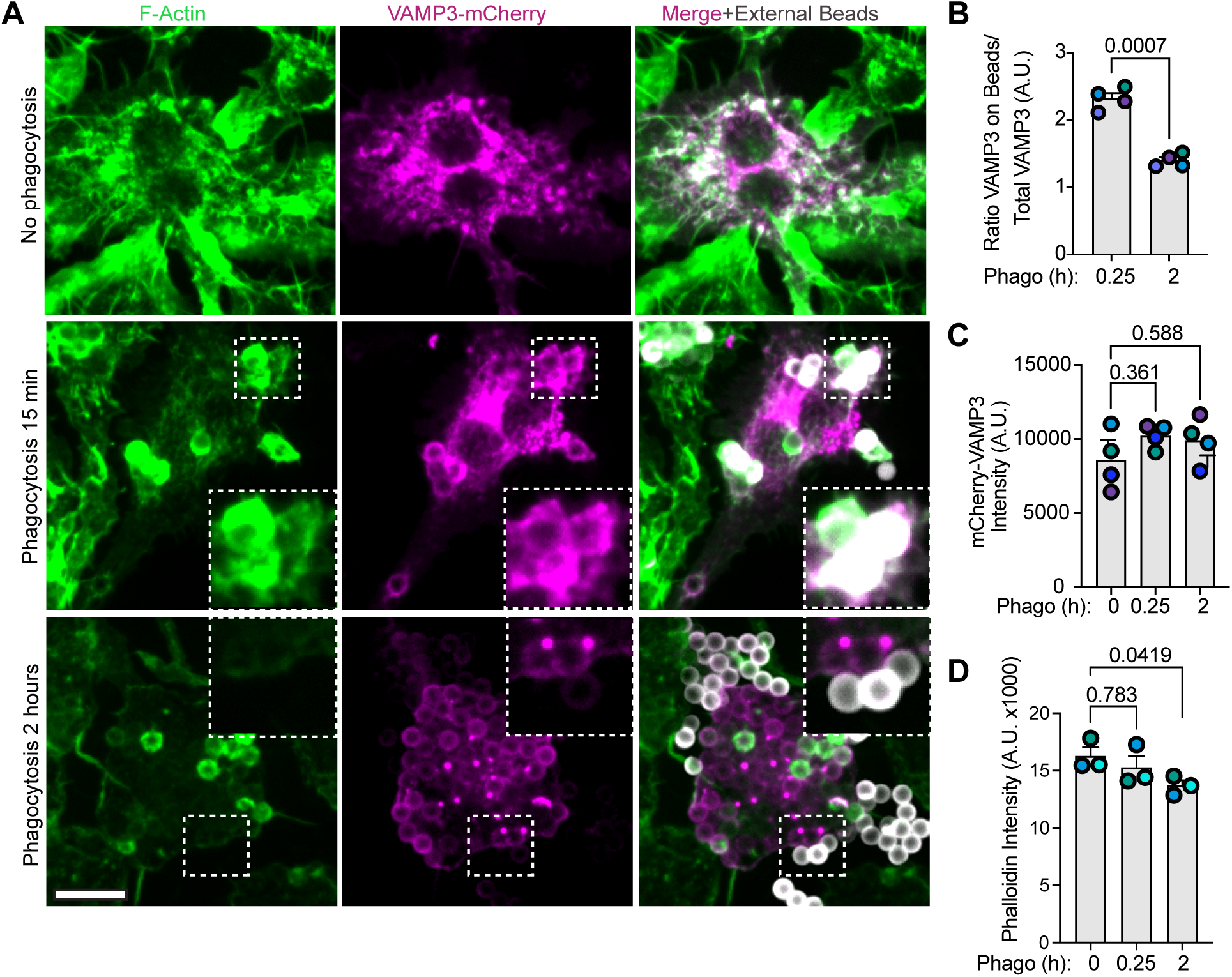
Localized exocytosis onto phagocytic cups fails in exhausted macrophages. **A.** RAW macrophages overexpressing mCherry-VAMP3 (magenta) engulfed IgG-opsonized 3 µm beads for 0, 15 min or 2 h. Cells were then fixed and labelled for F-actin (green) and external beads (white). Inserts are magnifications of bound external beads (dashed boxes). Scale bar, 10 µm. **B.** Quantification of the ratio of fluorescence intensity of VAMP3 on phagocytic cups relative to total VAMP3 fluorescence of that cell as shown in A. Data are mean ± SEM from 4 independent experiments with 75-150 cells per condition per replicate examined. Sample means were compared using two-tailed, paired Student’s t-test. **C, D.** Total fluorescence intensity of VAMP3 (C) and F-actin (D) per cell. Data are shown as mean ± SEM from at least 4 (C) or 3 (D) independent replicates. Sample means were compared using repeated measures, one-way ANOVA & Dunnett’s post-hoc test. p-values are shown.

**Figure 7:**
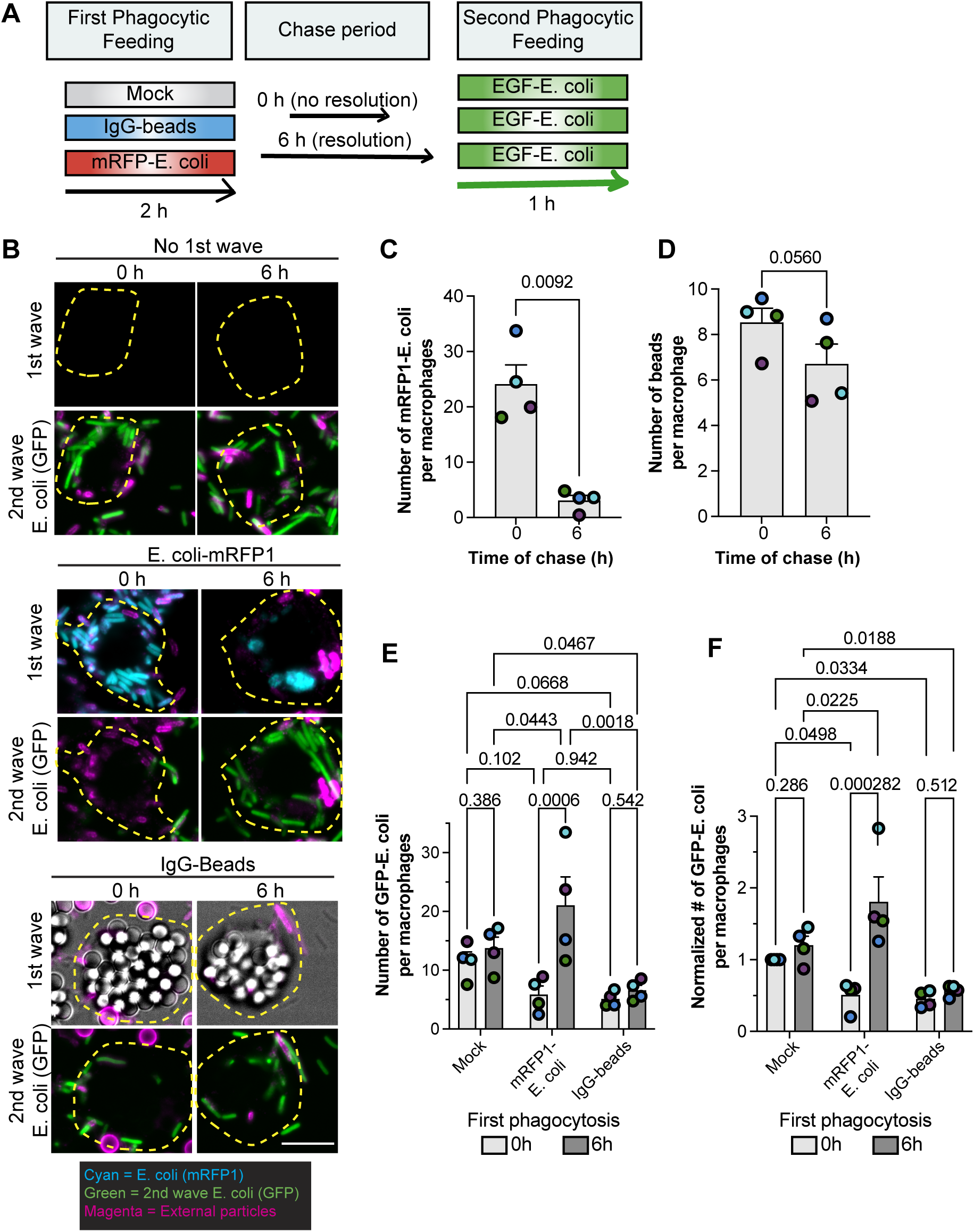
Phagosome resolution is required to restore phagocytic appetite. **A.** Schematic showing our strategy to test if phagosome resolution recovers appetite post phagocytic exhaustion. Briefly, macrophages were fed a first round of phagocytosis with IgG-beads, mRFP1-E. coli, or no particles (mock), followed by either no chase (no resolution) or 6 h to allow phagosome resolution, before being allowed to engulf GFP-*E. coli* for 1 h (second feeding). **B.** Using the strategy described in A, shown are images of RAW cells that engulfed GFP-*E. coli* (second feeding; green) after mock phagocytosis, or engulfment of *E.coli*-mRFP1 (cyan), or IgG-beads (grayscale) with no chase (0 h) or phagosome resolution (6 h). Magenta indicates external particles and dashed lines delineate cell boundaries. Scale bar: 10 µm. **C, D.** Quantification of mRFP1-labeled *E. coli* (C) or IgG-beads (D) at the conclusion of the first phagocytic feeding (0 h) or after phagosome resolution (6 h); phagosomes with bacteria, but not with beads, were resolved after 6 h. Data was analysed by a paired, two-tailed Student’s t-test. **E, F**. Quantification of phagosomes with GFP-*E. coli* (second feeding) in macrophages that were mock-fed, or engulfed mRFP1-E. coli, or IgG-beads with no resolution time (0 h) or after 6 h post-initial feeding. Data represented as mean ± SEM from N=4 independent trials, with each sample and experiment based on 120-300 cells. For E, data was analysed by repeated-measures, two-way ANOVA and Tukey’s post-hoc test. For F, data was tested with repeated-measures, two-way ANOVA and Dunnett’s Multiple Comparison test. p-values are disclosed.

To further test if phagosome resolution is a prerequisite for macrophages to recover their phagocytic appetite, we performed the two-phagocytic feeding assay in macrophages exposed to ikarugamycin starting at 2 h after the first phagocytic feeding; ikarugamycin is a clathrin inhibitor that arrests phagosome resolution (Lancaster et al., 2021). We note that the final readout of this experiment (the number of GFP-*E. coli*) likely varies due to the compounding heterogeneity caused by differences in the first phagocytic feeding, rates in phagosome resolution and recycling, and the effect of clathrin inhibition of biosynthetic pathways (Duncan, 2022). We mitigated this by using two different time points for resolution (4 and 6 h) and normalizing data. As before, we observed that macrophages that were fed consecutive rounds of phagocytosis (no time for resolution) displayed reduced uptake of GFP-*E. coli* (Fig. 8A, B; first two columns). In comparison, those macrophages that were given 6 h to resolve phagosomes from the prior engulfment, recovered their phagocytic appetite (Fig. 8A, B, column 2 vs. column 8). Treatment of ikarugamycin alone for 4 or 6 h did reduce uptake (Fig. 8A, B, column 1 vs. columns 5 and 9). However, we note that the combined effect of prior phagocytosis and 4 h of ikarugamycin was significantly more inhibitory then 4 h of ikarugamycin alone (column 5 vs. 6) and recovery of phagocytosis after 6 h of chase was blunted if cells were exposed to ikarugamycin (columns 8 vs 10; columns 4 vs 6 at 4 h, trended but did not reach statistical significance with p ∼ 0.1). These observations suggest that phagosome resolution helps macrophages recover their phagocytic appetite after phagocytic exhaustion. Since resolution reforms lysosomes (Lancaster et al., 2021), we attribute this to recovery of membrane reservoirs.

**Figure 8:**
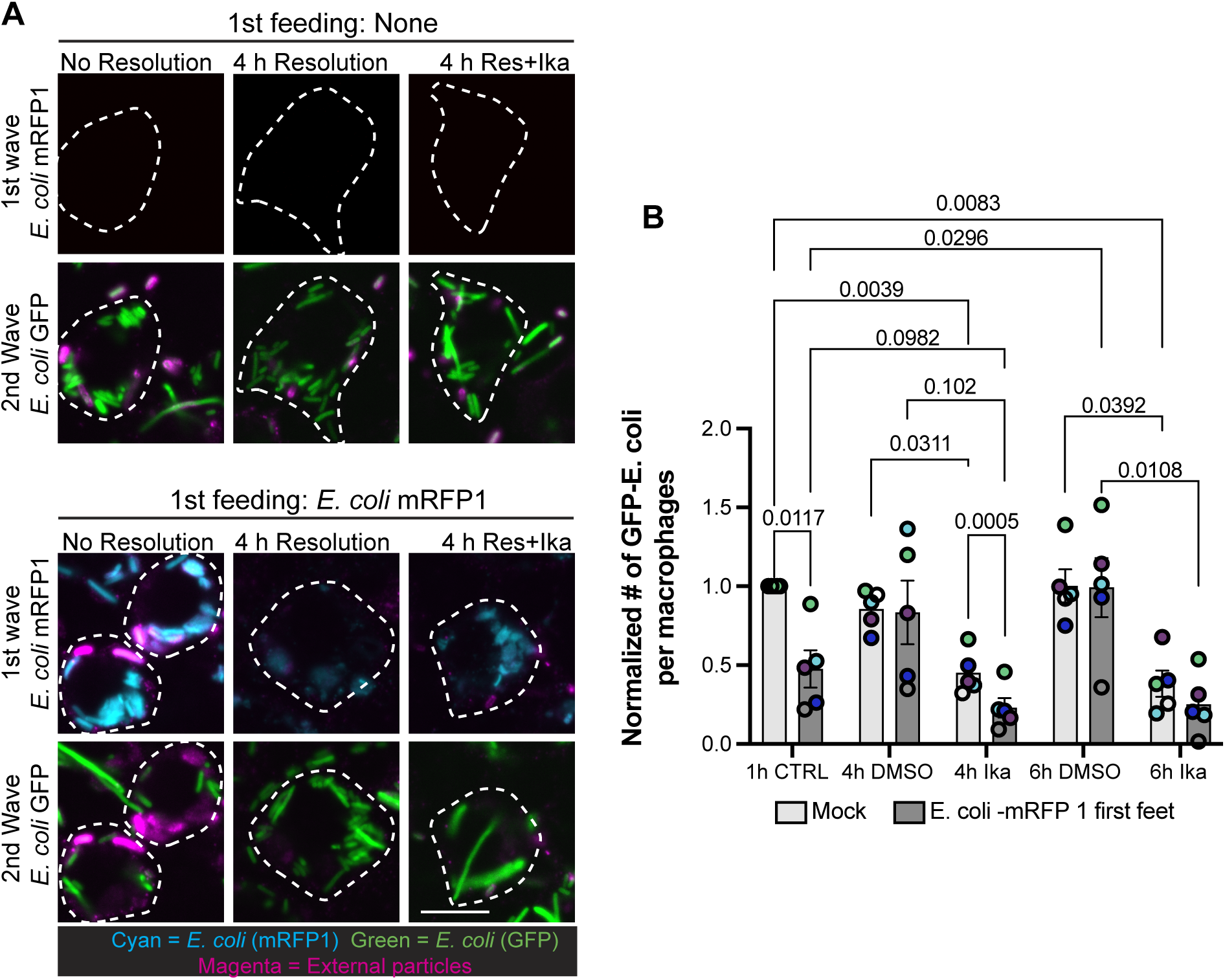
Inhibition of phagosome resolution with clathrin inhibitors impairs appetite recovery post-phagocytic exhaustion. **A.** Micrographs of RAW cells that engulfed GFP-*E. coli* (second feeding; green) after mock phagocytosis or engulfment of *E.coli*-mRFP1 (cyan) for 2 h, followed by resolution (0 h chase), or 4 h (or 6 h, not shown) with and without 10 µM ikarugamycin, which was added at 2 h post-phagocytosis to allow phagosome maturation but interfere with phagosome resolution. External particles were immunolabeled with anti-*E. coli* antibodies (magenta). Dashed lines indicate cell boundaries. Scale bar: 10 µm. **B.** Normalized number of GFP-*E. coli* engulfed per macrophage. Treatment groups and chase periods are as described in A. Data are shown as normalized mean (to mock first feeding, vehicle, and no resolution) ± SEM from N=5 independent experiments, where 100-200 cells were counted per condition per experiment. Data were compared by repeated-measures two-way ANOVA and Tukey’s post-hoc test. P values for statistically significant (p<0.05) comparisons shown. Specifically, there is a drop in phagocytosis after first feed (column 1 vs. column 2). This is recovered after 6 h of resolution (column 2 vs. column 8) but inhibited if ikarugamycin is included (column 8 vs. column 10). Ikarugamycin alone does reduce phagocytosis (column 1 vs. columns 5 and 9), but ikarugamycin combined with prior phagocytosis has stronger inhibitory effect than ikarugamycin alone at 4 h of resolution (column 5 and 6).

### Evidence for multiple processes imparting phagocytic appetite exhaustion

Our work mostly focused on using IgG-opsonized 3 µm beads or using *E. coli.* However, the types of particles and engaged phagocytic receptor may lead to deviations from our observations described above. In fact, below is evidence that phagocytic appetite exhaustion and recovery is complex and subject to multiple mechanisms. To illustrate, if phagocytic exhaustion depended solely on macrophages running out of membrane reservoirs, then one would expect that macrophages would engulf the same total amount of membrane surface area of particles/phagosomes. To test this, we fed IgG-opsonized 1.1, 3, or 6.1 µm diameter-beads to macrophages until they reached capacity. Cells were then labelled with a cytosolic dye to better define the phagosomes within cells by exclusion of the dye (Sup. Fig. S5A). We then estimated the number of particles engulfed, which not surprisingly was greater for smaller particles than larger ones (Sup. Fig S5B). However, when we calculated the total surface area and volume of all these phagosomes, we observed a massive difference between 1.1 and 6.1 µm beads. Macrophages that engulfed 6.1 µm beads could ingest ∼500-1000x more surface area and volume than macrophages that engulfed 1.1 µm beads (Sup. Fig S5C, D). Additionally, cells that engulfed 6.1 µm beads appeared to enlarge in their volume as indicated by orthogonal views of macrophages (Sup. Fig. S5A); while we refrain from ascribing actual values due to light refraction by these large particles, these observations suggest that particle properties may impact macrophage volume at capacity as described in Fig. 3. More intriguingly, this implies that macrophages that engulfed 1.1 µm beads had more space and area available to them, but instead they stopped engulfing these smaller beads before reaching that limit. Collectively, we submit that other mechanisms exist that determine the phagocytic exhaustion other than depletion of endomembranes and that their relative contribution may depend on particle properties such as size.

## Discussion

Over the years, we have learned many details about the processes required for phagocytosis and phagosome maturation (Fountain et al., 2021; Freeman and Grinstein, 2014; Lancaster et al., 2019; Richards and Endres, 2017). Surprisingly though, and while it is obvious that macrophages must have a phagocytic capacity limit, there is a remarkable dearth of knowledge on the mechanisms encoding this limit, an issue previously highlighted by Zent and Elliott (Zent and Elliott, 2017). Like these authors, we posited that such mechanisms could include phagocytic receptor depletion, negative feedback signaling, membrane depletion, and/or physical crowding caused by the presence of many phagosomes. These mechanisms are non-mutually exclusive, and their individual level of contribution may vary with the type of phagocytosis, phagocyte, and target particle. Our data evince that membrane reservoir depletion is a major limiting factor of phagocytic appetite in macrophages at capacity during FcγR-mediated phagocytosis using 3 µm beads.

### Surface levels of Fcγ receptors drop in exhausted macrophages, but does not appear to cause phagocytic exhaustion

Phagocytic receptors that engage ligands on target particles are entrapped within the phagosome (Booth et al., 2002; Lin et al., 2016); this should lead to a reduction in receptor levels at the surface as they are consumed with each nascent phagosome. In fact, Pinney *et al*. demonstrated that FcγR levels dropped during antibody-dependent cell phagocytosis of tumour cells, leading to “hypophagia”, *i.e.,* phagocytic exhaustion (Pinney et al., 2020). We also observed a significant decline in surface levels of CD16A and CD64 using microscopy and flow cytometry in macrophages fed IgG-coated 3 µm beads to capacity. While we were concerned that externally bound particles occluded access of staining antibodies to surface receptors, we still observed a reduction in surface receptor staining in cells fed IgG-coated sRBCs even when the external sRBCs were lysed into “loose” membrane ghosts, we do not think this is due to occlusion of antibody binding to receptors. Despite these observations, we argue that Fcγ receptors levels do not limit appetite as macrophages approach their capacity. If this was the case, macrophages overexpressing FcγRIIa should have increased their phagocytic capacity, but they did not. Thus, while surface levels of Fcγ receptors ostensibly drop in exhausted macrophages, we think this is not a limiting factor for at least this phagocytic model. Additionally, receptor properties such as receptor clustering, dynamics, and signaling capability may be different between resting and exhausted macrophages (Freeman et al., 2018; Kern et al., 2021; Lin et al., 2016; Zhang et al., 2010); this should be explored in future work.

### Exhausted macrophages do not shrink in their cell size but lose surface membrane folds

Focal exocytosis of endosomes and lysosomes is proposed to help form phagocytic cups and replace the plasma membrane consumed during phagocytosis, thus preventing the dwindling of phagocyte size and surface area (Bajno et al., 2000; Lee et al., 2007; Samie et al., 2013). However, to the best of our knowledge, it was not known if macrophages that engulfed to their full capacity would maintain their size or shrink as they consumed to their maximal capacity. We found that macrophages at capacity do not shrink – instead macrophages either retain their size, as estimated when engulfing beads < 3.1 µm in diameter, or increase their cell size when macrophages engulf to capacity 6.1 µm beads or yeast conidia (Fig. 3 and Sup. Fig. S5). Nevertheless, we perceived the surface membrane topology of exhausted macrophages to be distinct from resting macrophages, lacking most membrane folds and extensions, which is consistent with past observations using guinea pig macrophages (Petty et al., 1981). Thus, we surmise that surface folds are consumed to accommodate phagosomes to capacity, but cells stop short of reducing their size. Consistent with this, Gβ4-deleted phagocytes had extensive membrane ruffles compared to wild-type cells, which boosted their plasma membrane surface area and their total phagocytic capacity (Winer et al., 2024).

Interestingly, when using 3 µm beads for phagocytosis, “full” macrophages displayed a raspberry-like morphology because the packed phagosomes bulged out. We initially postulated that this meant that the membrane was under tension, collapsing membrane folds like a “tense balloon”. However, neither the membrane tension probe, FLIPPER®, nor AFM measurements supported this notion. Instead, surface tension may be abated, driven partly by a drop in cortical actin in exhausted macrophages. Such a phenomenon would not only collapse the surface membrane folds but would also cause the plasma membrane to contour the underlying phagosomes like a wet blanket, while only forming shallow phagocytic cups on remaining externally bound particles. It will be interesting to understand the signals that disturb the cortical actin, which may intersect with Rho-family GTPases and phosphoinositide signaling (Araki et al., 1996; Botelho et al., 2000; Caron and Hall, 1998; Marshall et al., 2001).

### Membrane reservoir depletion is a key determinant of phagocytic exhaustion

Phagocytosis is aided by local exocytosis of endosomes and lysosomes, depending on the size of the particle (Bajno et al., 2000; Lee et al., 2007; Samie et al., 2013; Vinet et al., 2008). Nevertheless, it was not known if macrophage exhaustion occurred before depletion of these membrane reservoirs. We show here that macrophages at capacity suffer a large loss of “free” lysosomes, which we previously observed (Lancaster et al., 2021). This was not surprising given that phagolysosomes are the end stage of maturation, prior to resolution. Notwithstanding this, early phagosome maturation is thought to be transient and thus, one would anticipate that endosomes reform even as macrophages reach capacity; thus *a priori*, endosomes should not be depleted in these macrophages. However, imaging and biochemical data show that a VAMP3-labelled pool of endosomes is significantly tied up as phagosomes in exhausted macrophages. This suggests that a major determinant of phagosome exhaustion is the depletion of internal membrane pools and impaired exocytic delivery of membranes to form phagocytic cups. To support this, phagosome resolution, which reforms lysosomes, was a prerequisite to recover phagocytic appetite. Feeding indigestible beads or blocking phagosome resolution of digestible cargo prevented macrophages from recovering their appetite.

Overall, we propose that depletion of endo-lysosomal membrane reservoirs is a major determinant of phagocytic exhaustion. Of course, this process may not universally apply to all forms of phagocytosis as suggested by a distinct total phagosomal volume and membrane area occupied by macrophages that ingested 6 µm, 3 µm, or 1 µm beads. For example, it is possible that other sources of membrane like the Golgi apparatus and/or the endoplasmic reticulum contribute to the maximal uptake of larger particles. Moreover, it is likely that additional, concurrent mechanisms are at play such as a collapse of the actin cortex, possibly by negative feedback signaling, and/or metabolic changes that could pause phagocytic appetite until particles are digested. Future work will be required to understand the cortical actin collapse and whether receptor dynamics and signaling play a major role in phagocytic exhaustion and recovery.

### Physiological processes that may lead to phagocytic capacity and appetite exhaustion

In our study, we challenged macrophages to continuously engulf particulates for 2-3 hours leading to phagocytic exhaustion. A key question is whether there are analogous physiological challenges that could lead to phagocytic exhaustion *in vivo*. To the best of our knowledge this has not been deliberately investigated, but we anticipate that physiological challenges that could lead to phagocytic exhaustion include localized bacterial infections, biofilms, incessant clearance of senescent erythrocytes in the spleen by red pulp macrophages, tissue damage causing high apoptotic load, and antibody-dependent cell-mediated cytotoxicity used to treat tumours. In fact, Grandjean *et al*. observed macrophages eating multiple tumours cells after anti-CD20 injection and Kwiencinshi *et al*. saw dense *S. aureus* bacterial communities during skin infections that may lead to phagocyte exhaustion (Grandjean et al., 2021; Kwiecinski et al., 2021). Additionally, Kupffer macrophages exhibit phagocytic exhaustion (hypophagia) after engulfing B cells decorated with anti-CD20 antibodies *in vivo*, though it is unclear if these cells had achieved phagocytic maximal load (Pinney et al., 2020). Thus, whether phagocytes engulf to their maximal theoretical capacity or manage to avoid this *in vivo* is a separate and interesting question to study. For example, phagosomes containing degradable particles will undergo resolution and flux fast enough to avoid attaining their theoretical maximal phagocytic capacity. Thus, there is a need to explore phagocytic capacity limits and appetite exhaustion in *in vivo* with methods like intravital imaging.

## Methods and Materials

### Cell line culture and transfection

RAW 264.7 murine macrophages (ATCC TIB-71; American Type Culture Collection, Manassas, VA) were cultured at 37°C and 5% CO_2_ in DMEM supplemented with 10% heat-inactivated FBS (Wisent Inc., Saint-Jean-Baptiste, QC) and tested for *Mycoplasma* routinely. RAW cells were passaged every 2-3 days by mechanically scraping the monolayer to dislodge the cells, followed by pipetting up and down to break up clumps. RAW cells were discarded after 20 passages. Prior to experimentation, cells were seeded at 30% seeding density on methanol-sterilized coverslips. RAW cells stably expressing Dectin-1 were used for phagocytosis of yeast. These cells were kindly provided by Dr. Nicolas Touret (University of Alberta) and were previously described (Lipinski et al., 2013). RAW^Dectin1^ cells were grown to full confluency in Permanox dishes (Nalge Nunc International Rochester, NY) that were previously coated for 1 h at room temperature with 6 mg/mL human fibronectin diluted in PBS (ThermoFisher Scientific).

Transfection of RAW cells with plasmid DNA was done with FuGENE HD (Promega). Briefly, 3 µL of FuGENE HD was mixed with 1 µg of plasmid in serum-free DMEM for 15 min at room temperature, for a final concentration of 6% v/v FuGENE HD and 20 µg/mL of plasmid. The plasmid:FuGENE solution was then diluted with DMEM complete media to a final concentration of 1 µg/mL of plasmid. RAW macrophages were transfected the day after seeding for 24 h at 37°C and 5% CO_2_, after which transfection reagents were removed with a PBS wash and replaced with complete DMEM medium. Macrophages were then incubated at 37°C and 5% CO_2_ overnight before beginning phagocytosis assays. The plasmid encoding VAMP3-mCherry was kindly provided by G. van der Bogaart (Radbound University Medical Center, Nijmegen, Netherlands; Addgene #92423) and described (Verboogen et al., 2017). The plasmid encoding EGFP-CAAX was kindly provided by Lei Lu (Nanyang Technological University, Singapore; Addgene # 86056) and described in (Madugula and Lu, 2016). The plasmid expressing FcγRIIa-GFP was obtained from the Grinstein lab (Booth et al., 2002).

### Primary macrophages and ethics statement

Mice were used following institutional ethics requirements under the animal user permit approved by the St. Michael’s Hospital Animal Care Committee, which is certified by the Canadian Council of Animal Care and the Ontario Ministry of Agriculture, Food, and Rural Affairs. Briefly, mice were anesthetized with 5% isoflurane administered by inhalation, followed by cervical dislocation before limb bone dissection to obtain bone marrow. No experiments were performed on live animals.

Bone marrow-derived macrophages (BMDMs) were harvested from wild-type 7-9-week-old male and female C57BL/6J mice (Charles River Canada, Montreal, QC) as previously described (Hipolito et al., 2019; Weischenfeldt and Porse, 2008). Briefly, bone marrow was isolated from femurs and tibias through perfusion with phosphate-buffered saline (PBS) using a 27G syringe. RBCs were lysed using a hypoosmotic treatment. For BMDMs, cells were plated according to experimental requirements in DMEM supplemented with 10% fetal bovine serum, 20 ng/ml recombinant mouse macrophage colony-stimulating factor (Gibco, Burlington, ON), and penicillin/streptomycin antibiotics. Media was changed every 2 days. Experiments were conducted on days 7–9.

### Bacterial and fungal strains and growth

*E. coli* DH5α was transformed with the pBAD::mRFP1 plasmid, whichS was kindly provided by Robert Campbell, Michael Davidson, and Roger Tsien (University of California at San Diego, La Jolla, CA, USA, Addgene, plasmid # 54667) and described in (Campbell et al., 2002), or with pZsGreen vector (catalog no. 632446; Takara Bio USA, Inc., Ann Arbor, MI). *E. coli* strains were grown overnight at 37°C on Luria-Bertani (LB) agar plates, and colonies were subsequently cultured overnight at 37°C in LB broth under agitation. Agar- and broth media were supplemented with 100 μg/ml ampicillin (BioShop Canada Inc., Burlington, ON). For *E. coli* expressing ZsGreen, 1% D-glucose (BioShop) was also added to the LB plates and overnight culture broth, but not the subculture broth, to suppress leaky expression of ZsGreen from the lac operon. Otherwise, overnight *E. coli* cultures were sub-cultured at 1:100 dilution and grown at 37 °C until mid-log phase. At this point, 5 mM L-arabinose (BioShop) or 1 mM IPTG (Millipore-Sigma, Burlington, ON) were added to respectively induce expression of mRFP1 and ZsGreen expression for 2-3 h. After induction, *E. coli* was washed 3 times in PBS and then immediately used live in phagocytic assays or fixed in 4% paraformaldehyde (PFA; Electron Microscopy Sciences, Hatfield, PA) in PBS and stored for a few weeks.

Filamentous targets of *Legionella pneumophila* (*L. pneumophila*) were obtained as previously described (Prashar et al., 2012; Prashar et al., 2013). Briefly, *L. pneumophila* Lp02 strain (mRP1-expressing *Legionella*) from glycerol stocks were streaked onto Buffered Charcoal-Yeast Extract (BCYE) agar and grown at 37°C and 5% CO_2_. After 72-96 h, colonies were harvested and cultured by triplicate for 24 h at 37°C in Buffered Yeast Extract (BYE) media at 100 rpm. Afterwards, bacteria cultures were diluted to an OD_600_ of 0.05 and cultured for another 16–18 h or until the cultures reached an OD_600_ of 3.5–4.0 (Molofsky et al., 2005). Cultures enriched in long filamentous bacteria (>100 µm) were centrifuged at 1000 x*g* for 15 min, washed twice with 1x PBS, and fixed at room temperature in 15 mL of 4% PFA for 20 min on a rotator. Fixed mRFP-*Legionella* filamentous targets were stored in 4% PFA at 4°C until use. The *Saccharomyces cerevisiae* (yeast) INVSc1 strain was grown as described by the manufacturer (ThermoFisher Scientific). Overnight liquid cultures were harvested by centrifugation at 1000 *xg* for 10 min, washed twice with PBS, and fixed at room temperature with 4% PFA for 20 min. The fixed targets were stored at 4 °C until use. *Aspergillus fumigatus* (UAMH 2978) was grown on potato dextrose agar incubated at 15°C for 7 days to generate conidia, which were then collected in PBS + 0.05% Tween80, followed by centrifugation at 10,000 xg, washed twice with PBS, and fixed at room temperature with 4% PFA for 20 min. The fixed targets were stored at 4 °C until use.

### Phagocytic Particle Preparation

To prepare IgG-opsonized plain polystyrene beads, 1.1, 3, or 6.1 μm beads (Bangs Laboratories and Sigma-Aldrich) were opsonized with 4 mg/ml of human IgG (I8640 or I4506; Sigma-Aldrich) for 1 h at room temperature. Beads were then washed and resuspended in PBS. For the preparation of IgG-coated sheep red blood cells (sRBCs; Innovative Research, Novi, MI), a stock solution was prepared at 1.92×10^6^ sRBCs/µL in PBS. We then opsonized 96×10^7^ sRBCs with 0.4 mg/mL anti-sheep RBCs (FisherScientific) in a total volume of 500 µL PBS at room temperature in a rotator. After 1 h, IgG-coated sRBC were washed 3 times in PBS (300 xg, 2 min), and stored at 4°C in PBS until use. After a week, the remaining opsonized sRBCs were thrown away and a fresh stock solution was prepared.

*E. coli* bacteria were grown as above and given to macrophages unopsonized. PFA-fixed mRFP-*Legionella* filaments were washed three times with PBS by centrifugation at 10,000 x*g*. Then, bacterial filaments were opsonized by resuspending the bacterial pellet in a 4 mg/ml solution of human IgG (Sigma-Aldrich) in PBS and allowing the sample to mix on a rotator overnight at 4 °C. Opsonized *Legionella* were washed three times as described above and resuspended in PBS.

For yeast targets, yeast cells were washed 3 times with PBS to remove residual PFA, resuspended in culture media and presented to the macrophages at a ratio of 25:1 target/cell. The cells were incubated at 37 °C for 2 h, after which they were processed for transmission electron microscopy as described below. For the preparation of IgG-opsonized *Aspergillus fumigatus* conidia fixed in 4% PFA, conidia were washed 3 times with PBS to remove residual PFA, resuspended in culture media and presented to the macrophages at a ratio of 100:1 target/cell. The cells were incubated at 37 °C for 2 h, after which they were processed for confocal microscopy imaging.

For the preparation of opsonized magnetic beads, 3 µm COMPEL COOH modified magnetic beads (Bangs Laboratories) were first crosslinked to bovine serum album (BSA). The COMPEL beads were washed twice in activation buffer consisting of filter-sterilized 100 mM 4-Morpholineethanesulfonic acid (MES) (Biobasic, Canada), pH 4.5, using a magnet to separate beads out of suspension after each wash. Beads were then prepared as 1% w/v suspension in 100 mg/mL of filter sterilized 1-Ethyl-3-(3 -dimethylaminopropyl)carbodiimide-HCl (EDAC; E2247, Sigma-Aldrich) in activation buffer, and incubated for 15 min while rotating at room temperature. The beads were then washed twice with Coupling buffer (filter sterilized 100 mM MES, pH 7.4) and resuspended to 1% w/v in filter sterilized 2% w/v BSA solution in Coupling buffer and incubated for 2 h while rotated at room temperature. Uncoupled or singly bound EDAC was quenched by washing the beads twice with quenching solution (filter sterilized 100 mM MES, pH 7.4, containing 40 mM glycine and 1% w/v Normal goat serum), resuspended to 1% w/v in quenching solution, and incubated for 30 min while rotated at room temperature. Beads were then washed twice with storage buffer consisting of PBS containing 1% w/v Normal goat serum. To opsonize the beads with antibodies, the BSA-coupled magnetic beads were suspended to 1% w/v in storage buffer containing 1:100 mouse anti-BSA antibody (Sigma, B2901) and incubated on rotation for 60 min at room temperature. The beads were washed 3 times with storage buffer then resuspended to 2% w/v in storage buffer.

### Phagocytic assays

Phagocytosis was accomplished with the following target to macrophage ratios that dependent on the extent of phagosome completion and exhaustion: 20–400 *E. coli* rods per RAW cell; 10 3.87-μm beads per cell, and 25–100 1.1, 3.0, or 6.1 μm beads per RAW cell; and 50 IgG-sRBCs per macrophage. Target attachment was synchronized by spinning cells at 300 × g for 5 min at 4°C, followed by phagocytosis between 15-min to 3 hours at 37°C. Unbound targets were then washed off with PBS. As required, cells were further incubated at 37°C for indicated times to allow phagosome maturation before processing for microscopy, Western Blotting, or scanning electron microscopy.

If live bacteria were presented to macrophages, 200 µg/mL of gentamicin (Gibco) was applied to the cells for 20 min post-phagocytosis and after washing unbound bacteria. For assays using two rounds of phagocytosis, the procedure for both the first and second rounds of phagocytosis was the same as described above, except the secondary phagocytic challenge was done after the first round of phagocytosis at specified times. The secondary challenge was then followed with no chase or 1-h chase to elicit further maturation before further processing for microscopy or lysate preparation.

For the synchronized phagocytosis of mRFP-*Legionella* filamentous targets, macrophages were cooled down to 15 °C for 10 min and subsequently challenged with opsonized bacterial targets at a ratio of 150:1 target/cell. Afterwards, cells were centrifuged at 15 °C and 300 *xg* for 5 min and subsequently incubated for 15 min at 37 °C to favour bacteria binding. Fresh medium was then given to cells and incubated for another 75 min at 37 °C. At the end of the incubation period, cells were fixed with 4% PFA in PBS and further processed for microscopy.

For phagocytosis of yeast and *A. fumigatus* conidia, macrophages were cooled down to 15 °C for 10 min and subsequently challenged with targets at rations described in sections above. Afterwards, cells were centrifuged at 15 °C and 300 *xg* for 5 min. Fresh medium was then given to cells and incubated for 2 hr at 37 °C. At the end of the incubation period, cells were fixed with 4% PFA in PBS and further processed for microscopy.

### Pharmacological inhibitors

Inhibitors and vehicle controls (typically, dimethyl sulfoxide (BioShop)) were applied after phagocytosis and maintained until fixation or the conclusion of the experiment. For clathrin inhibitors, cells were incubated with 0.5–2.0 μg/ml ikarugamycin (Sigma-Aldrich) 40 min to 2 h after the start of phagocytosis and continued for up to 4 additional hours.

### Fluorescent cell labelling and immunocytochemistry

For cytoplasm labeling prior to phagocytosis, cells were washed with PBS then incubated with 5 μM Carboxy Fluoroscein Succinimidyl Ester (CFSE, Invitrogen) in PBS for 15 min at 37°C. CFSE solution was then removed and replaced with complete DMEM medium, incubated for 5 min, before removing the media and replacing with fresh complete DMEM.

For immunocytochemistry, cells were fixed at the conclusion of the experiment with 4% PFA, followed by 100 mM glycine for 15 min to quench unreacted PFA. To identify extracellular targets after phagocytosis, secondary anti-human IgG antibodies or anti-rabbit IgG antibodies at 1:1000 dilution or rabbit anti-*E. coli* antibodies (Bio-Rad Laboratories) at 1:100 dilution in 2% BSA in PBS were applied to fixed, but non-permeabilized cells for 30 min. To stain all particles, cells were then permeabilized with 0.1% Triton-X100 in PBS for 10 min, followed by 30 min blocking with 2% BSA in PBS before antibody incubation.

For staining of surface CD16 by indirect immunofluorescence and for microscopy, after fixation and blocking with 5% skim milk, cells were incubated with rabbit anti-CD16a/ CD16b polyclonal antibody (1:1000, cat. # BS-6028R, ThermoFisher Scientific) for 1 h, then washed with PBS and incubated for 1 h with a secondary donkey-anti rabbit antibodies conjugated to DyLight 488nm (Novus Biologicals, Toronto, ON). Cells were then washed three times with PBS and mounted using DAKO fluorescence mounting media (Agilent, Mississauga, ON).

For immunolabeling of endogenous LAMP1 proteins, cells were fixed in 4% PFA for 15 min at room temperature, permeabilized with methanol at −20°C for 10 min, followed by 30 min of blocking with 2.5% BSA diluted in PBS. Afterwards, cells were incubated with a 1:100 dilution of rat anti-mouse LAMP1 antibodies (clone 1D4B, Developmental Studies Hybridoma Bank) for 1 h at 37 °C, washed three times with PBS, followed by incubation for 1 h at room temperature with 1:1000 fluorescently tagged secondary antibodies, and subjected to another round of washes with PBS. Cells were mounted in Dako fluorescent mounting medium (Agilent Technologies, Inc.).

### Viability and mitochondrial activity of exhausted macrophages

To assess cell viability, cells were seeded onto an 18 mm glass coverslip and then subjected to phagocytosis (100 targets per cell), mock treatment, or a 2 min treatment with hydrogen peroxide (positive control). Cells were then washed with PBS and incubated with a 10,000x dilution of Sytox^TM^ Deep Red (ThermoFisher) diluted in serum-free DMEM for 15 min at 37°C, followed by 3 washes with PBS, and then imaged live to score the number of Sytox-positive cells over total cells. To determine the activity of mitochondria, control and exhausted macrophages were labelled with 0.4 µM MitoTracker Red (Thermofisher) and 0.2 µM MitoTracker Green FM (Thermofisher) in serum-free media for 30 min at 37°C. Cells were then imaged live and analysed by Mander’s co-localization. Briefly, we scored the proportion of active mitochondria (MitoTracker Red) within the total mitochondria population (MitoTracker Green) by sampling the Mander’s co-localization of red in the green channel.

### Fluorescence microscopy

Confocal images were acquired by a Quorum Diskovery spinning-disk confocal microscope system (Quorum Technologies, Inc.) equipped with an inverted fluorescence microscope (DMi8; Leica) using either an Andor Zyla 4.2-megapixel scientific complementary metal–oxide– semiconductor camera or an Andor iXON 897 EM-CCD camera (Oxford Instruments). Alternatively, a Quorum Wave FX-X1 spinning-disk confocal microscope system (Quorum Technologies, Inc.) was used equipped with an inverted fluorescence microscope (DMI6000B; Leica Microsystems) using either a Hamamatsu ORCA-R2 camera (C10600-10B) or a Hamamatsu ImagEM Enhanced EM charge-coupled device (CCD) camera (Hamamatsu Corporation). Images were acquired using a 63× oil immersion objective (1.4 NA) or a 40× oil immersion objective (1.3 NA). For live-cell imaging, coverslips were enclosed in a Leiden chamber and immersed in DMEM supplemented with 10% FBS and mounted in a microscope-mounted environmental chamber maintained at 37°C and 5% CO_2_.

### Fluorescence Image Analysis

Image processing and quantitative analysis were performed using Fiji (Schindelin et al., 2012) or Volocity (Quorum). Image enhancements were completed without altering the quantitative relationship between image elements. Figures were prepared with Adobe Illustrator (Adobe, Lehi, UT).

For quantification of phagocytic index using beads, non-externally labelled beads were counted and marked manually per macrophage. For *E. coli*, if images were a z-stack, these were first collapsed using maximum intensity method to reconstruct a 2D image. Then phagosomes were counted manually as above OR semi-manually as follows: a mask was first created to remove *external E. coli,* then mRFP1 signal expressed by *E. coli* was thresholded, followed by watershed to separate distinct bacteria. Particle count was then applied for objects over 10 pixel^2^ to exclude noise.

To quantify CD16A/B cell surface levels, z-stack images were collapsed using sum of intensities method. Then fluorescence intensity values were acquired for all complete cells in a field of view and normalized to control condition.

For the quantification of macrophage surface area and volume, image stacks of macrophages ectopically expressing eGFP-CAAX were reconstructed in Volocity, then processed to fill in holes within the 3D-reconstructed objects to assess total volume of the objects. 3D-reconstructed objects were then thresholded by size, with cells being classified as having a total volume > 100 µm^3^. Cells that were touching one another were separated manually by encircling a cell with an ROI, then clipping the 3D-reconstructed object to within the ROI.

For the quantification of the number of free lysosomes in each cell, we used Volocity to separate touching objects in the LAMP1 channel using an object size guide of 0.29 μm^3^ (determined by assessing endosome and lysosome size in resting macrophages). LAMP1-positive objects were considered free lysosomes if their volume was >0.02 μm^3^ but <5 μm^3^ and they were not touching filament-containing phagolysosomes (applied a mask to the filament or beads). For endosomes, phagosome-associated VAMP3-mCherry signal contouring or proximal to beads was manually removed after background correction. Images were then thresholded to create binary images of the samples, and the binary images were processed by object segmentation using Fiji’s Watershed function. Events above 0.038 μm^2^ were counted using Fiji’s Analyze Particles function. Large objects that were erroneously segmented were corrected by subtracting the number of additional erroneous segments from the total particles detected. The treatments were finally normalized to the no-phagocytosis group, which was considered to contain 100% free lysosomes or endosomes.

To quantify VAMP3-mCherry and phalloidin-stained F-actin intensities in whole cell and in nascent phagocytic cups, z-stacks were collapsed using the *sum slices* method to reconstruct a 2D image in FIJI. The total intensity for each protein was then extracted per cell. Then, a mask was created using the external beads and applied to the different channels to measure the fluorescence intensity on or proximal to the external beads.

### Flow cytometry and analysis

To label external CD16 and CD64 in live cells for flow cytometry analysis, 900,000 RAW264.7 cells were seeded 18-24 hours before phagocytosis. Following phagocytic uptake with 3 µm beads or sRBCs, cells were suspended in blocking media consisting of RPMI 1640 Medium (ATCC Modification, Gibco) supplemented with 10% FBS. Cells were blocked for 10 min on ice then centrifuged at 300 x*g* for 5 min. The cell pellet was resuspended in 100 µL blocking media containing anti-mouse CD16 antibodies conjugated to FITC (Cat #158008, BioLegend, CA) and anti-mouse CD64 antibodies conjugated to PE/Cyanine7 (Cat # 139314, BioLegend) at 1:100 dilution. Cells were incubated for 30 min on ice in the dark. Propidium iodide (BioShop) was added at 1:3000 dilution immediately before washing. Cells were washed twice by centrifugation at 300 xg for 5 min then resuspending the cell pellet in HBSS (Gibco) supplemented with 2% FBS. Samples were run on a Beckman Coulter CytoFLEX S flow cytometer, and analysis and mean peak values were calculated using the CytExpert Software.

### Fluorescence Lifetime Imaging Microscopy (FLIM)

To measure the plasma membrane tension, RAW macrophages were seeded in an open µ-slide with a glass bottom (Ibidi, Fitchburg, WI, USA). Cells were challenged or not with IgG-opsonized plain polystyrene beads as described above or treated for 10 min with a hypo-osmotic media (15% phenol-red free DMEM, 85% milliQ water). Cells were then incubated with the Flipper^®^ membrane probe (Cytoskeleton, Denver, CO, USA; (Colom et al., 2018)) for 15 min prior to live-cell imaging at 37°C and 5 % CO_2_. Cells were maintained at 37°C and 5% CO_2_ during imaging at the Advanced Optical Microscopy Facility (University Health Network, Toronto, ON, Canada) under a super-resolution microscope Leica DMi8 (Leica TCS SP8 X Scanner) equipped with a PicoHarp 300 TCSPC FLIM module. Cells were excited with a white light laser pulsed at 80 MHz and observed with a 93x HC PL APO CS2 objective (1.3 NA, Glycerol immersion). Leica Application Suite X (LAS X) v3.5.7 software was used for image acquisition and SymPhoTime 64 v2.3 software for FLIM analysis following manufacturer’s guidelines for Flipper^®^.

### Atomic Force Microscopy (AFM)

To measure the Young’s elastic modulus of the cell, AFM measurements were performed using a JPK NanoWizard 4 AFM system (Bruker, Billerica, MA, USA). Cells were kept at 37°C using the AFM BioCell and in phenol-red free media supplemented with 20 mM HEPES to maintain pH levels at ambient CO_2_. Quadratic pyramidal cantilever probes (MLCT-BIO, Bruker) with a spring constant of 0.1 Nm^−1^ and a resonating frequency of 38 kHz were used to perform indentation force–distance measurements. All tips were calibrated in liquid with contact-based and contact-free methods according to the manufacturer’s instructions. Set point (2.4 nN), z-length (2 µm), and extend speed (5 µm s^−1^) were kept constant. Per each cell and condition, measurements of the nucleus and the cytoplasm near or away from beads were performed, where 20 force curves per position were collected. To do this, 20 x 20 µm images were generated by the conversion of probe deflection upon indentation into a z-length (height through quantitative imaging (QI) Mode with the same settings previously described). QI mode is forced-based imaging resulting in high-resolution mapping of samples; each pixel is associated with a force curve. QI imaging permits one to define regions to measure or avoid measuring (e.g. beads for example). Data were processed using JPK Data Processing Software (Bruker, USA) by fitting the resulting force curves using an established Hertz model describing the deformation of two perfectly homogenous surfaces. This is estimated by expressing the normal force F_n_ of the probe with a radius of R as: 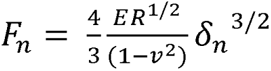 where δ_n_ is the indentation depth perpendicular to the sample and ν is the Poisson’s ratio of the sample (Fujii and Okajima, 2019). Mean values were calculated from all processed measurements (excluding beads) and plotted as mean Young’s elastic modulus in kilopascals (kPa).

### Transmission and Scanning Electron Microscopy

For scanning electron microscopy, samples were fixed in EM fixative (2.5% glutaraldehyde and 2.5% formaldehyde in 0.1 M sodium cacodylate buffer, pH 7.3) for at least 2 h. Fixed samples were processed by SickKids Nanoscale Biomedical Imaging Facility (The Hospital for Sick Children, Toronto, ON, Canada): samples were rinsed with 0.1 M sodium cacodylate buffer (0.1 M Sodium Cacodylate, pH 7.3, 0.2 M sucrose) and post-fixed with 1% osmium tetroxide for 1.5 h. After post-fixation treatment, samples were rinsed again with 0.1 M sodium cacodylate buffer and dehydrated with a series of ethanol solutions of increasing ethanol percentage from 70-100%. Samples were then dried with a CO_2_ critical point dryer and coated with a thin layer of gold by sputter coating. Samples were imaged on a Hitachi FlexSEM 1000 II Scanning Electron Microscope (Hitachi High-Tech, Inc.) in secondary electron acquisition mode with electron beam voltage set to 7.0 kV at SickKids Nanoscale Biomedical Imaging Facility (The Hospital for Sick Children).

For transmission electron microscopy, samples were fixed in with 2.5% glutaraldehyde and 2% PFA in 0.1 M sodium cacodylate buffer, pH 7.3 for 2 h at room temperature. Fixed samples were processed by SickKids Nanoscale Biomedical Imaging Facility (The Hospital for Sick Children). Briefly, samples were rinsed with 0.1 M sodium cacodylate buffer, post-fixed with 1% osmium tetroxide and 1.25% potassium ferrocyanide in cacodylate buffer for 1.5 h at room temperature. After post-fixation treatment, samples were rinsed again with 0.1 M sodium cacodylate buffer, then rinsed twice with distilled water (dH2O), and stained for 30 minutes in 4% uranyl acetate aqueous solution, followed by two additional dH2O rinses. Samples were dehydrated with increasing ethanol concentrations from 70-100% and infiltrated with Epon resin (SPI-Pon 812 Embedding Kit, Structure Probe, Inc. West Chester) at 1:3, 1:1 and then 3:1 Epon:ethanol ratios for 1.5 h each, in sequence, followed by incubation in 100% Epon for 3 h.

For polymerization, the Permanox dishes were filled with 3 mL 100% Epon and placed in at 70 °C overnight. Samples were then separated from the dishes and 80 nm thin sections were cut and placed on copper grids. Samples on grids were then stained with uranyl acetate and lead citrate as described elsewhere (Tapia et al., 2012). Samples were imaged using a Hitachi H7500 TEM Transmission Electron Microscope (Hitachi High-Tech, Inc.) with electron beam voltage set to 80kV, captured with Megaview II Camera (iTEM software) at the University of Toronto Scarborough.

### Electron microscopy analysis

For the quantification of SEM populations, cells were qualitatively classified into one of three subpopulations of membrane ruffles. Cells that appeared to have large, sharp membrane sheets and protrusions on their surface that extended well beyond the base surface of the cell were classified as “High Ruffling”. Cells that appeared to have more shallow protrusions, such as “veins” on the surface or shallow pits, that covered at least 50 percent of the visible cell surface were classified as “Medium Ruffling”. Conversely, cells that displayed protrusions on less than 50 percent of their visible surface, or cells that displayed almost no protrusions, were classified as “Low Ruffling”. For each sample, the percent of cells associated with each subpopulation was calculated across all images in a sample.

We quantified endosome and lysosome-like organelles using ultrastructural classification of late endosome, lysosome, and autophagic compartments was described here (Hess and Huber, 2021). Briefly, single 80 nm ultrathin sections were analyzed at low magnification and >ten macrophages were assessed per condition, per trial. Only macrophages sectioned at a plane through approximately the middle of the nucleus were chosen for further analysis. Single membranous compartments that resembled endosomes and lysosomes as defined above were identified throughout the whole cell section at a magnification of 5,000x, after which the total cell bodies were imaged at 20,000x to allow for a better visualization of endosomes and lysosomes.

### Cell Lysates and Membrane Fractionation

Cell lysates were prepared post-experimental treatment by washing 3x with ice-cold PBS and then incubating with 2x Laemmli Sample Buffer (0.5M Tris, pH 6.8, glycerol and 10% sodium dodecyl sulfate (SDS)). Samples were then scraped to ensure full release from the substratum and lysates passed through a 27-gauge needle ten times to break up genomic DNA. To each tube, 10% ß-mercaptoethanol and 5% bromophenol blue was added. Finally, lysates were boiled at 65°C for 15 min.

For isolation of phagosomes containing magnetic beads, cells were first washed twice with ice-cold PBS, then incubated in ice-cold homogenization buffer (250 mM sucrose, 10 mM CaCl_2_, 1:50 protease inhibitor cocktail (P8340, Sigma) and PhosStop phosphatase inhibitor (1 tablet/10 mL; Roche, Mississauga, ON) for 5 min. While on ice, cells were dislodged from the surface by scraping, then lysed by passing the cell suspension 20 times through a 27-gauge needle. The lysate was then incubated for 5 min on ice. Magnetic phagosomes were separated from the remaining lysate by magnetic attraction. Phagosomes were washed twice with PBS containing 1% normal goat serum and supplemented with 1:50 protease inhibitor cocktail, using the magnet to pull the beads out of suspension at the end of each wash. The magnetic phagosomes were lysed with 2x Laemmli containing 10% β-mercaptoethanol for Western Blot analysis. Laemmli buffer was added to the remaining cell lysates (left over after phagosome pulldown) to a final 1x concentration.

### Sodium dodecyl sulphate (SDS)-polyacrylamide gel electrophoresis (PAGE) and Western blotting

Cell lysates and membrane fractions were resolved in home-made SDS-PAGE gels ranging from 8% to 18% acrylamide or using 4-20% gradient mini-PROTEAN TGX precast gels (BioRad, Mississauga, ON). Electrophoresis was done in Tris-glycine running buffer (25 mM Tris-HCl, 250 mM glycine, 1% SDS). Proteins were then transferred onto polyvinylidene difluoride membranes. Membranes were blocked with 5% BSA in Tris-buffered saline (TBS) containing 0.1% Tween-20 (TBS-T) for 1 h at room temperature, then incubated overnight at 4°C with the following primary antibodies diluted in 1% BSA in TBS-T: 1:1000 rat anti-LAMP1 monoclonal antibodies (clone number: 1D4B, Santa Cruz Biotechnology) or 1:1000 rabbit anti-LAMP1 monoclonal antibodies (clone number: C54H11, Cell Signalling Technology), 1:400 rabbit anti-Cellubrevin/VAMP3 polyclonal antibodies (104103(SY), Synaptic Systems), 1:1000 rabbit anti-CD16A/B polyclonal antibodies (BS-6028R, ThermoFisher Scientific), 1:1000 rabbit anti-GAPDH antibodies (clone number: 14C10; Cell Signaling Technology). HRP-conjugated donkey anti-rabbit or anti-rat antibodies (Bethyl Laboratories and Cell Signalling Technology) were used at 1:10,000 in 1% BSA with TBS-T for 3 h at room temperature. Streptavidin-HRP (Cat. 3999S, Cell Signaling Technology) was incubated at 1:1000 for 1 h. Membranes were washed three times with TBS-T for 5 min each wash and developed with Clarity Western ECL Substrate (Bio-Rad Laboratories) or Immobilon Crescendo Western HRP Substrate (WBLUR0500, Millipore). Blot images were acquired using the ChemiDoc Imaging System (Bio-Rad Laboratories) and analyzed by ImageLab (Bio-Rad Laboratories). The background was automatically determined and subtracted from each band by the software, which employed a rolling disk method with a disk size of 10 mm to determine the background intensities along the lane. The background values corresponding to the positions of the bands were used to subtract the background from the respective bands.

### Statistical Analysis

Data are presented as the mean ± SEM for experiments where a sample mean was acquired from at least three independent experiments. For Western blotting, data is shown as mean ± STD. The number of cells assessed in each experiment is indicated within the figure legends. Statistical analysis was performed using GraphPad Prism software (GraphPad Software, Inc.). Unless data is normalized to a control, data were assumed to be normally distributed. For statistical testing between two conditions, a paired, two-tailed Student’s t-test or a Mann-Whitney test was used; if data was normalized, then a one-sample t-test was used. For statistical analysis of data from multiple groups with one variable, we used a repeated-measures one-way ANOVA coupled to a Tukey’s or Dunnett’s multiple comparisons test. If missing values/unequal sample size existed between groups, a mixed-effects test was applied, followed by a Dunnett’s test. If comparing multiple groups and two parameters, then we used a repeated measures two-way ANOVA coupled to Tukey’s or Sidak’s multiple comparisons test. Typically, Greisser-Greenhouse correction was applied. Friedman’s test was used if data were non-parametric (*e.g.* control was normalized) followed by Dunn’s post-hoc test. Typically, p < 0.05 was considered significant, but we opted to disclose actual p values.

## Acknowledgements

We would like to thank the Electron Microscopy Facility at the Hospital for Sick Children in Toronto, the Core Facility Imaging Centre at St. Michael’s Hospital, the Imaging Facility CAMiLoD at the University of Toronto, and the Centre for the Neurobiology of Stress (CNS) at the University of Toronto Scarborough for their invaluable support and advice.

## Abbreviations

AFM: Atomic Force Microscopy
BSA: Bovine serum albumin
*E. coli*: Escherichia coli
DMEM: Dulbecco’s Modified Eagle’s Medium
FBS: Fetal Bovine Serum
FcγR: Fcγ receptor
FLIM: Fluorescence Lifetime Imaging Microscopy
GADPH: Glyceraldehyde-3-Phosphate Dehydrogenase
GFP: Green Fluorescent Protein
IgG: Immunoglobulin G
LAMP1: Lysosome-Associated Membrane Protein 1
mRFP1: Monomeric Red Fluorescent Protein 1
PBS: Phosphate-buffered Saline
PFA: paraformaldehyde
RAW cells: RAW 264.7 macrophages
sRBC: sheep red blood cell
VAMP3: Vesicle-Associated Membrane Protein 3

---

**Supplemental Figure S1: Exhausted macrophages are viable and have active mitochondria relative to naïve macrophages.** RAW cells were exhausted with either PFA-fixed *E. coli*-mRFP1 or -iRFP1, or 3 µm polystyrene beads. **A.** Control and exhausted macrophages were stained with Sytox^TM^ Deep Red staining. Cells were also treated with 4% hydrogen peroxide for 2 min as a positive control. Dashed lines exemplify cell boundaries. Scale bar: 10 µm. **B**. Percentage of cells with Sytox^TM^ Deep Red nuclear staining from treatments in A. Data are shown as means ± SEM from N = 3 independent experiments, where at least 250 cells were assessed per condition per experiment. **C**. Micrographs of RAW cells after mock phagocytosis or 2 h phagocytosis with PFA-fixed *E. coli*-iRFP (blue pseudocolour) or 3 µm polystyrene beads. MitoTracker Red accumulation in mitochondria indicates active mitochondria while MitoTracker Green stains the whole mitochondria population. Dashed lines exemplify cell boundaries. Scale bar: 5 µm. **D**. Proportion of active mitochondria stained as described in *C*. Data are mean ± SEM of 30 cells from N=3 independent experiments. Data were compared by repeated-measures one-way ANOVA and Dunnett’s post-hoc test, p values are shown.

**Supplemental Figure S2: Representative flow cytometry plots of CD16A and CD64 in macrophages before and after phagocytic exhaustion.** RAW macrophages were fed IgG-beads (A, B) or IgG-coated sRBCs (C, D) for 0 (black trace), 15 (green trace), or 120 min (magenta trace). After phagocytosis, external sRBCs were lysed with hypotonic shock. Unpermeabilized cells were then stained with co-stained FITC-labelled anti-CD16A (A, C) or PC7-labelled anti-CD64 (B, D). Surface fluorescence was quantified by flow cytometry as per Methods. Quantification is shown in Figure 2.

**Supplemental Figure S3: Reduction in endo-lysosomal like organelles in exhausted RAW and primary macrophages. A.** Naïve RAW cells expressing VAMP3-mCherry (top) or after engulfing filamentous *Legionella* (grayscale). Dashed lines show the outline of macrophages. Scale bar = 10 µm. **B.** Representative electron micrographs of naïve RAW^Dectin1^ macrophages (left) or after 2 h of phagocytosis of yeast (darker structures within macrophages). Yellow arrows indicate organelles that resemble endosomes/lysosomes/autophagosomes. Scale bar = 500 nm. **C.** Number of endosome-lysosome-like structures per macrophage. Each data point is the number of yeast visible in an 80 nm ultrathin section, with 41 and 26 resting and exhausted macrophages scrutinized from N=2 experiments (blue dots from experiment 1, pink from experiment 2). **D**: BMDMs from male (squares) and female (circles) mice were given no particles (Mock phagocytosis, lanes 1-3) or allowed to engulf IgG-opsonized magnetic beads (Magnetic beads, lanes 4-6) for 2 h. Phagosomes were then magnetically isolated and analyzed with Western Blot. LAMP1 and VAMP3 abundance were probed as a proxy for lysosome and endosome content, respectively. GAPDH was used as a loading control in whole cell lysates (WCL) and post-magnetic fraction (PML) is content remaining in lysate after magnetic separation; Magnetic phagosome fraction is represented by M. **E, G**: Quantification of percent LAMP1 (E) and VAMP3 (G) remaining in lysate after magnetic separation. Percent remaining protein is defined as the percent ratio of GAPDH-normalized protein content in the PML fraction (lanes 2 and 5 for mock phagocytosis and magnetic beads, respectively) to GAPDH-normalized protein content in the WCL fraction (lanes 1 and 4 for mock phagocytosis and magnetic beads, respectively). **F, H**: Analysis in E and G, respectively, but normalized to mock phagocytosis to account for experimental variation in absolute values. Data represented as mean ± STD of N=3 independent experiments. Sample means were compared using two-tailed, paired Student’s t-test. *: p<0.05; **: p<0.01 (E, H) or with the one sample t-test (F, I).

**Supplemental Figure S4: Cortical F-actin is abated in exhausted primary macrophages. A, C:** BMDMs derived from male and female mice were fed IgG-coated 3 µm beads (A) or IgG-opsonized sRBCs (C) for 0, 15, or 120 min and then fixed and stained with phalloidin (magenta, F-actin) and DAPI (nuclei, cyan). sRBCs were also stained with anti-rabbit antibodies (C, grayscale). Scale bar = 10 µm. **B, D**: F-actin levels based on normalized phalloidin fluorescence from z-stacks collapsed by sum-intensity per cell from N=5 independent experiments using 10 images (3-8 cells per image) per sex per condition per experiment. Circle and square icons respectively represent means acquired from female and male mice. Data are represented as mean ± SEM and were analysed by the Friedman test (B) or Kruskal-Wallis test (D, difference is due to a missing sample for the 15-min condition) and Dunn’s post-hoc test; p-values are indicated.

**Supplemental Figure S5: Particle size may affect the mechanism of phagocytic exhaustion. A.** Representative orthogonal confocal planes of RAW macrophages at rest or after 3 h of engulfing 1.1 or 6.15 µm IgG-coated beads. Z-stacks were acquired from top to bottom of cells. Shown at the X-Y plane is a slice near the site of cell attachment to the coverslip. YZ and XZ orthogonal views are shown corresponding to the yellow lines. Scale bar = 10 µm. **B.** Estimated number of beads in macrophages after 3 h of phagocytosis. **C, D**. Total surface area (C) and total volume (D) of all phagosomes in cells at capacity. This was estimated based on the total phagosomes in *A* and the radius of each bead using surface area and volume formula for a sphere. Data are shown as the mean ±SEM of N=3 independent experiments based on 30 cells per condition per experiment. Sample means were statistically tested by repeated measures one-way ANOVA and the post-hoc Tukey’s test. p values are disclosed.

