## Supplementary figures and images for "Depletion of endomembrane reservoirs drives phagocytic appetite exhaustion in macrophages"

### Supplemental Figures S1, S2, S3, S4, and S5

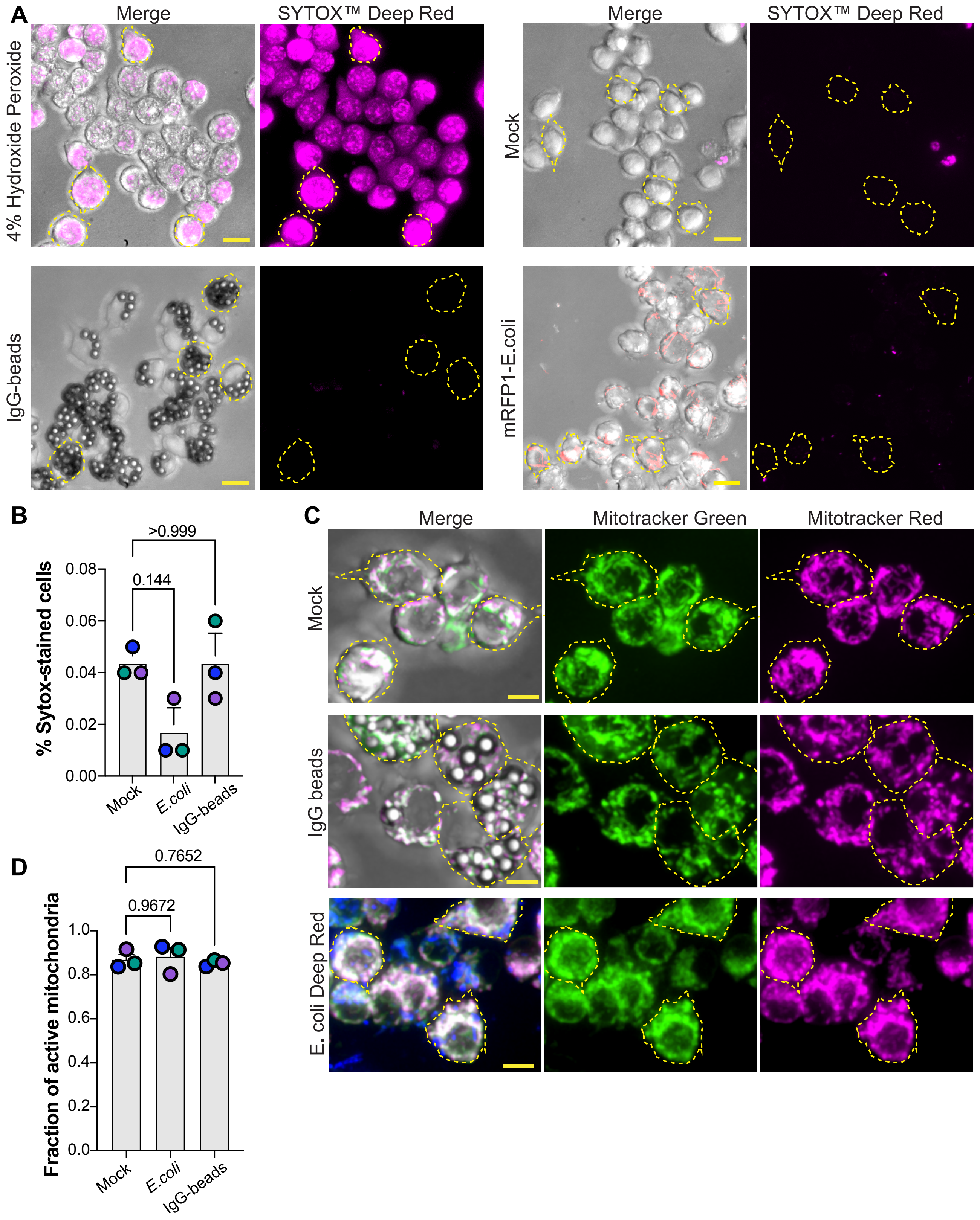


Fountain et al., Sup. Fig. S1

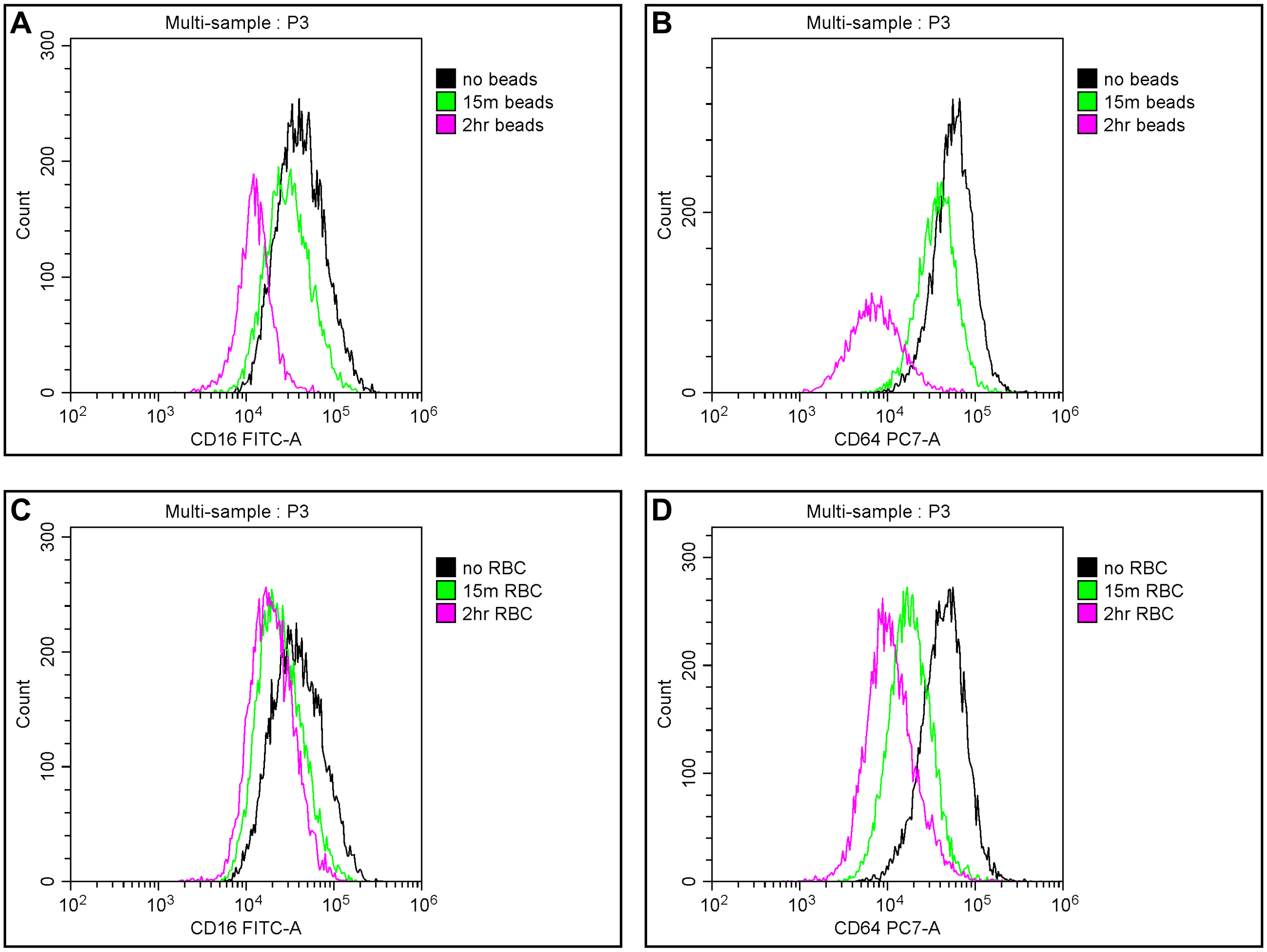


Supplemental Figure S2

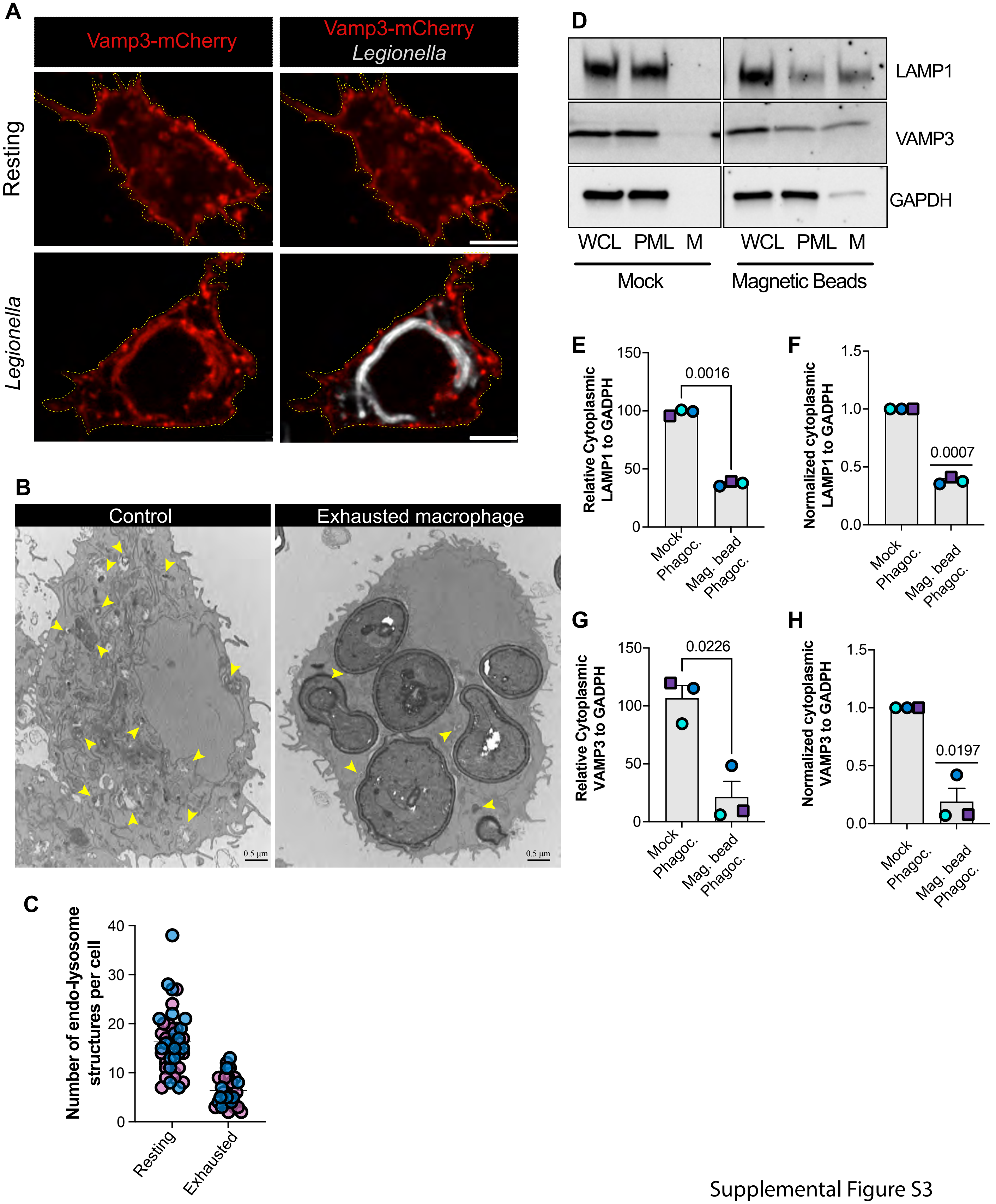


Supplemental Figure S3

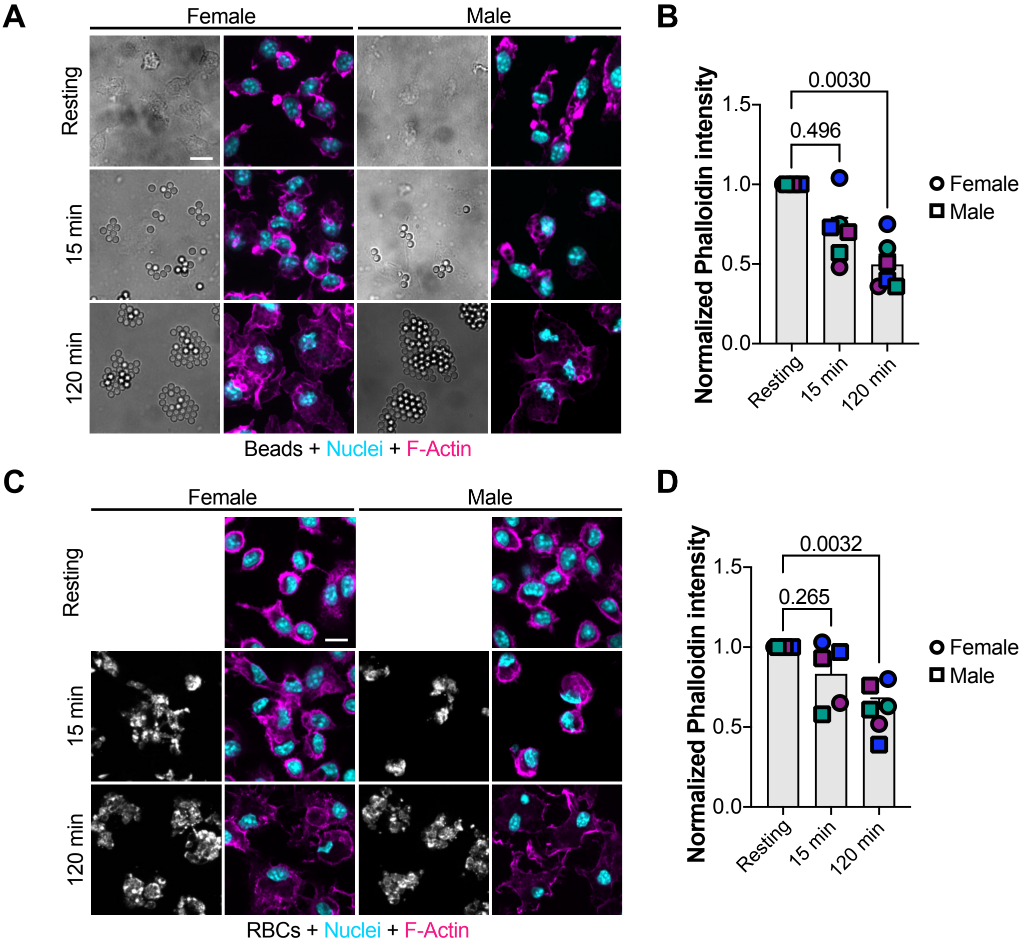


Supplemental Figure S4

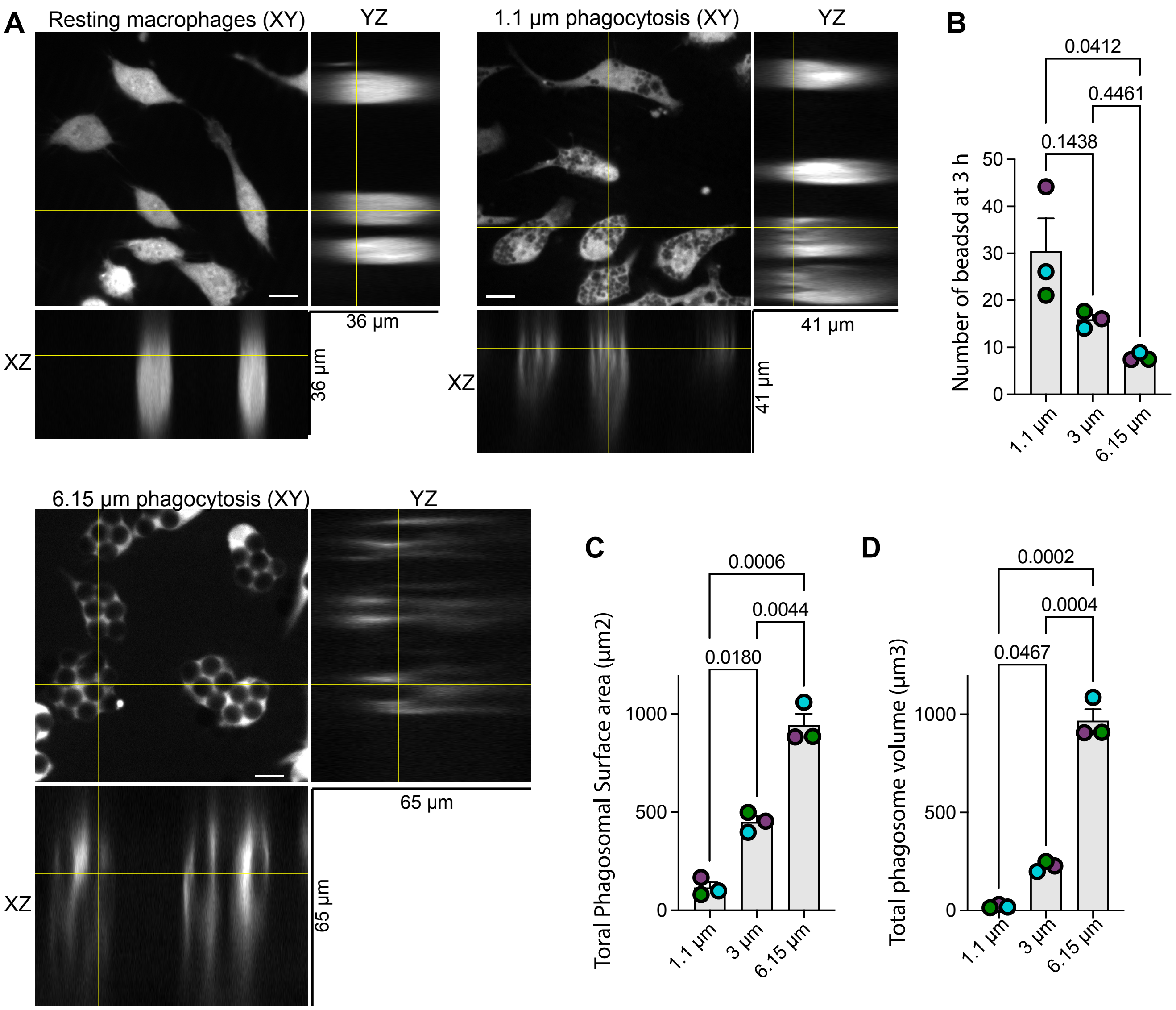
